# Activating an Interleukin-4-FLT3-STAT6 axis in Multipotent Progenitors Restores Lymphopoiesis in Inflammation and Aging

**DOI:** 10.64898/2025.12.19.695491

**Authors:** Jingfei Yao, Yuting Wang, Yi Zhang

## Abstract

Chronic inflammation and aging skew hematopoiesis toward myelopoiesis at the expense of lymphoid output. We screened type 2 and anti-inflammatory cytokines to identify extrinsic signals capable of restoring lymphoid lineage commitment in hematopoietic stem and progenitor cells (HSPCs). Interleukin-4 (IL-4) specifically inhibited inflammation-induced myelopoiesis and shifted multipotent progenitor (MPP) differentiation toward the lymphoid lineage. IL-4 activated a STAT6-dependent transcriptional program in MPPs, increasing expression of lymphoid-specific genes. Mechanistically, the receptor tyrosine kinase FLT3, highly expressed in MPPs, interacted with IL-4Rα to facilitate STAT6 activation. In vivo, IL-4 reversed inflammation-induced hematopoietic imbalance and accelerated lymphoid recovery. In aged mice, IL-4 administration shifted the MPP composition toward lymphoid bias and restored B and T lymphocyte output. Long-term IL-4 treatment of aged mice improved immune, metabolic, physical, and cognitive functions; these rejuvenating effects were recapitulated by transplantation of IL-4-treated HSPCs. Promoting IL-4 signaling on MPPs may enable correcting hematopoietic dysregulation in inflammatory and aging-related conditions.

## Introduction

Hematopoiesis, the process of blood generation, is fundamental for maintaining immune function and systemic homeostasis^1,2^. The delicate balance between myelopoiesis and lymphopoiesis is dynamically regulated in responding to the physiological and environmental stimuli such as infection, inflammation, or aging^3–7^. At the core of the regulation are the hematopoietic stem and progenitor cells (HSPCs), which serve as the source of all blood cell lineages^8,9^. HSPCs are hierarchically organized. At the apex of this hierarchy, long-term hematopoietic stem cells (LT-HSCs) ensure lifelong blood production. LT-HSCs first differentiate into short-term HSCs (ST-HSCs), also referred to as multipotent progenitor (MPP) 1, which then give rise to lineage-restricted MPP subsets^10,11^. These subsets exhibit distinct lineage biases: MPP2 primarily produces megakaryocytes and erythrocytes, MPP3 predominantly generates granulocytes and monocytes, while MPP4 favors the development of lymphoid cells, including B and T lymphocytes^10–12^. Recent advances in barcoding and single-cell technologies have revealed that MPPs are established in parallel with HSCs during fetal development. Rather than being solely downstream products of HSCs, embryonically established MPPs contribute substantially to lifelong lymphoid output, highlighting their role in serving as key initiators of hematopoiesis^13–15^. This paradigm shift has prompted a deeper investigation into how fate decisions are regulated across different subsets of HSPCs. These processes are governed by a dynamic interplay between intrinsic genetic programs and extrinsic signals derived from the bone marrow microenvironment^16^. Together, these regulatory mechanisms maintain hematopoietic homeostasis by ensuring both the continuous production of blood cells and an adaptive response to physiological demands or pathological challenges, such as infection or tissue injury^17,18^.

Cytokines are pivotal regulators of hematopoiesis, acting as important extrinsic signals that guide HSPC fate under steady-state and stress conditions^7,19^. Pro-inflammatory cytokines such as interleukin (IL) –1β and tumor necrosis factor-alpha (TNF-α) are potent inducers of myelopoiesis, driving granulocyte and monocyte production to reinforce innate immunity during infections and inflammatory responses^5,20,21^. These cytokines activate signaling cascades and epigenetic modifications in HSPCs, promoting myeloid lineage differentiation while inhibiting lymphoid commitment^20–22^. While these mechanisms ensure robust innate immunity, persistent inflammation or aging disrupts hematopoietic equilibrium, skewing differentiation toward myelopoiesis at the expense of lymphopoiesis^23–25^. This imbalance contributes to immune dysfunction, chronic inflammation, and increased susceptibility to infections and metabolic disorders^26–28^. Despite advances in understanding pro-inflammatory drivers of myelopoiesis, the mechanisms for restoring lymphoid commitment and achieving balanced hematopoiesis under inflammatory or aging conditions are not fully understood.

Type 2 immunity is a critical component of the immune system, primarily involved in responses to helminth infections, allergic inflammation, and tissue repair^29,30^. It is orchestrated by type 2 cytokines, including IL-4, IL-5, IL-9, IL-13, IL-25, and IL-33, which are predominantly produced by Th2 cells, group 2 innate lymphoid cells (ILC2s), and other immune cells^31,32^. These cytokines regulate the differentiation of Th2 cells, stimulate eosinophil and mast cell responses, and drive antibody class switching to immunoglobulin E (IgE), contributing to pathogen defense and hypersensitivity reactions^31,32^. The cytokines that drive type 2 immunity, such as IL-4 and IL-13, also exhibit significant anti-inflammatory properties, which are vital for maintaining immune homeostasis and limiting tissue damage caused by excessive immune activation^33–36^. However, the roles of type 2 and anti-inflammatory cytokine signaling in regulating hematopoiesis under inflammatory conditions—particularly in lymphoid cell fate commitment—are poorly defined.

Here, by screening type 2 cytokines as well as anti-inflammatory cytokines *in vitro*, we found only IL-4 could restore hematopoietic balance by promoting lymphopoiesis while inhibiting myelopoiesis in HSPCs. Through transcriptomic profiling and functional assays using both inflammation and aging models, we showed that IL-4 reprogramed MPPs—but not HSCs—toward a lymphoid-biased fate via STAT6-dependent transcriptional mechanisms. Further analysis revealed that FLT3, which is selectively expressed in MPPs, cooperated with the IL-4 signaling pathway to enhance STAT6-mediated lymphopoiesis. Notably, IL-4 administration reversed inflammation– and aging-associated myeloid bias, restores lymphoid output, and improved systemic immune and metabolic functions. These findings highlight IL-4 as a promising therapeutic candidate for correcting hematopoietic dysregulation in inflammatory and aging-related conditions.

## Results

### IL-4 promotes lymphopoiesis of hematopoietic stem and progenitor cells

To identify potential external factors that might promote lymphopoiesis and inhibit myelopoiesis, we first performed a screen using various type 2 cytokines as well as some anti-inflammatory cytokines, including IL-4, IL-5, IL-6, IL-9, IL-10, IL-13, IL-25, and IL-33. To simplify the process, we utilized a well-established *in vitro* myeloid cell differentiation assay by incubating isolated LT-HSCs with essential cytokines^20,21^. Consistent with previous reports, IL-1β, a well-known pro-inflammatory cytokine, greatly increased myeloid cell differentiation (**Figure 1A** and **Figure S1A**). Among the cytokines tested, only IL-4 significantly inhibited IL-1β-induced myelopoiesis (**Figure 1A**). This result demonstrated that IL-4 specifically antagonizes IL-1β-induced myelopoiesis compared to other type 2 cytokines or anti-inflammatory cytokines. Further analysis revealed that the inhibition by IL-4 was primarily due to the loss of myeloid progenitors (**Figure 1B** and **Figure S1A**), indicating that IL-4 may act on early stages of hematopoietic differentiation.

**Figure 1.**
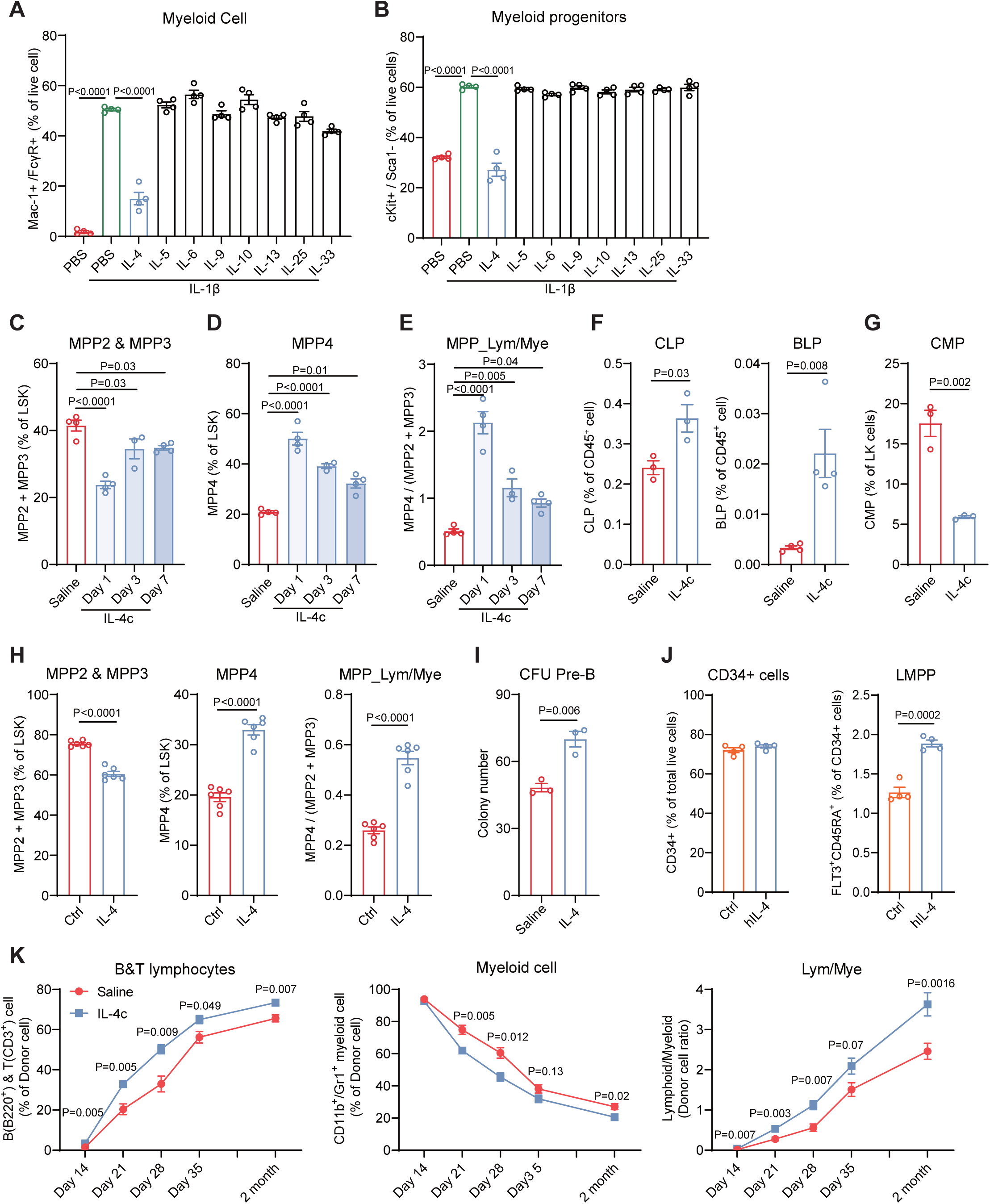
IL-4 promotes lymphopoiesis of hematopoietic stem and progenitor cells. **A-B**, *In vitro* myeloid differentiation of LT-HSC with or without 50 ng/ml IL-4. Results are shown as percentage of Mac-1^+^/FcyR^+^ mature myeloid cells (**A**) and c-Kit^+^/Sca-1^−^ myeloid progenitors (MyP) (**B**) upon 6-day culture (n=4). **C-D**, Percentage of BM MPP2 & MPP3 (**C**) and MPP4 (**D**) in HSPCs from WT mice with or without IL-4c injections (n=3-4). **E**, The ratio of BM MPP_Lym/Mye [MPP4/(MPP2&MPP3)] from WT mice with or without IL-4c injections (n=3-4). **F-G**, Percentage of CLP, BLP(**F**), and CMP(**G**) in CD45^+^ BM cells from WT mice with or without IL-4c injections (n=3-4). **H**, Percentage of MPP2 & MPP3, and MPP4 subpopulations within isolated HSPCs, and the MPP_Lym/Mye ratio, with or without 24 hours treatment using 50 ng/mL IL-4 *in vitro* (n=6). **I**, Pre-B colony forming unit (CFU) assays of IL-4c treated or untreated BM cells in methylcellulose after 7 days culture *in vitro* (n=3). **J**, Percentage of human CD34⁺ HSPCs and LMPPs after 24 h treatment with human IL-4 (n=4). **K**, Transplantation of HSPCs from IL-4c treated or untreated mice in lethally irradiated recipients, donor-derived lymphoid cells (**K, left**), myeloid cells (**K**, **middle**) and lymphoid-to-myeloid ratio (**K**, **right**) in peripheral blood at the indicated days post-transplantation (n=4). All data represent means ± s.e.m. Statistical significance was determined by one-way ANOVA with Tukey’s multiple-comparisons test (**A-E**) or unpaired two-tailed Student’s t-test (**F-K**).

To examine the role of IL-4 in early hematopoiesis *in vivo*, mice were injected with IL-4 complex (IL-4c; 2.5 µg IL-4 complexed with 12.5 µg anti–IL-4 antibody to enhance stability)^37^ for the indicated durations. Compared to saline-injected controls, IL-4c-treated mice showed a reduced percentage of LT-HSCs and ST-HSCs, while the total hematopoietic stem and progenitor cell (HSPC: Lin^−^Sca-1^+^c-Kit^+^) population remained unchanged (**Figure S1B** and **Figure S2A-S2D**). We then analyzed downstream lineage-restricted multipotent progenitor cell types, including MPP2, MPP3, and MPP4. IL-4c treatment decreased both the percentage and absolute numbers of MPP2 and MPP3, while increased that of the MPP4 (**Figure 1C-1D**, and **Figure S2E**), consistent with inhibited myelopoiesis observed *in vitro* (**Figure 1A-1B**). Since both MPP2 and MPP3 are myeloid-biased subsets distinct from the lymphoid-primed MPP4, we categorized MPP2 and MPP3 together as myeloid multipotent progenitors (MPP_Mye) and MPP4 as lymphoid multipotent progenitors (MPP_Lym). We also used the MPP_Lym/Mye ratio (MPP4/MPP2&MPP3) to reflect the pro-lymphoid shift in lineage commitment (**Figure 1E**). To exclude potential effects of the anti–IL-4 antibody presents in IL-4c, we separately injected IL-4 cytokine alone or anti–IL-4 antibody alone. IL-4 alone significantly increased the MPP_Lym/Mye ratio, but anti–IL-4 antibody had no effect (**Figure S2F**), confirming that the lineage-skewing effect of IL-4c is due to IL-4 itself rather than to the antibody component. Flow cytometry analysis further revealed changes in downstream progenitor populations following IL-4c treatment, including increased percentage of common lymphoid progenitors (CLP), and B-lymphocyte progenitors (BLP), concomitant with decreased percentage of common myeloid progenitors (CMP) (**Figure 1F-1G**). These changes were consistent with the observed shifts in multipotent progenitors, confirming that IL-4 promotes lymphoid progenitor expansion while reducing myeloid progenitor populations. We also analyzed other progenitors, including LK populations (Lin^−^Sca-1^−^c-Kit^+^), granulocyte-macrophage progenitors (GMP), megakaryocyte-erythrocyte progenitors (MEP) and megakaryocyte progenitors (MkP) (**Figure S2G-S2H**). Collectively, these results demonstrate the potent effect of IL-4 on shifting HSPCs differentiation towards lymphoid lineage.

Given that IL-4 regulates not only hematopoietic progenitors but also mature cell types *in vivo*, systemic IL-4 administration complicates the attribution of lineage bias specifically to HSPC differentiation. To study IL-4’s direct effects on HSPC fate, we cultured bone marrow-derived HSPCs *in vitro* with or without IL-4. We found that IL-4 treatment *in vitro* increased MPP4 population and decreased MPP2&3 population, but without affecting the total number of HSPCs (**Figure 1H** and **Figure S2I**). This rapid lineage reprogramming in a simplified culture system—free from confounding systemic signals—supports the hypothesis that IL-4 directly promotes a myeloid-to-lymphoid shift in HSPCs. To further examine whether an increased MPP_Lym/MPP_Mye ratio produces more lymphoid lineage cells, we performed pre-B colony formation assays in methylcellulose *in vitro*. We found that IL-4c-treated bone marrow produced more pre-B colonies than controls (**Figure 1I**). We next tested whether IL-4 influences human lymphoid-primed multipotent progenitors (LMPPs: CD34⁺CD38⁻CD90⁻CD45RA⁺FLT3⁺ cells). To this end, purified human bone marrow CD34⁺ HSPCs were cultured with recombinant hIL-4, which significantly increased the proportion of LMPP population without altering the overall frequency of CD34⁺ HSPCs (**Figure 1J**), consistent with the lineage bias observed in mouse HSPCs.

To directly evaluate IL-4’s effect on HSPC hematopoiesis *in vivo*, we transplanted HSPCs isolated from saline– or IL-4c-treated mice into recipient mice. Consistent with the increased MPP_Lym/MPP_Mye ratio and more pre-B colonies, IL-4c-treated HSPCs generated more mature lymphoid cells and fewer mature myeloid cells in peripheral blood (**Figure 1K** and **Figure S2J**). We also tested different doses of IL-4 treatment both *in vivo* and *in vitro*, and found that a low dosage of IL-4 is potent for inducing the MPP_Lym/Mye ratio without further improvement at higher doses (**Figure S2K**). Additionally, IL-4 treatment did not affect the viability of HSPCs (**Figure S2L**). These results collectively demonstrate that IL-4-induced HSPC differentiation shifts toward the lymphoid lineage, leading to predictable changes in peripheral blood cell composition.

### The IL-4-STAT6 signaling axis regulates genes important for lymphopoiesis

Next, we attempted to understand the molecular mechanism by which IL-4 shifts HSPC differentiation. Since previous studies have identified three pathways (STAT6, AKT, and ERK) that mediate IL-4-IL-4Rα signaling^38^, we used specific inhibitors to evaluate the contribution of each pathway. We found that only the STAT6 inhibitor completely blocked the IL-4-induced increase in the MPP4 percentage and MPP_Lym/MPP_Mye ratio, while the AKT and ERK inhibitors had no effect (**Figure 2A**). Consistently, IL-4 significantly activated STAT6 in HSPCs as indicated by the increased phosphorylated STAT6 level after 3 hours of treatment, and the activation persisted, albeit partially, even after 12 hours of treatment (**Figure 2B**). These results support that STAT6 is the primary downstream mediator of the IL-4’s effects on HSPCs.

**Figure 2.**
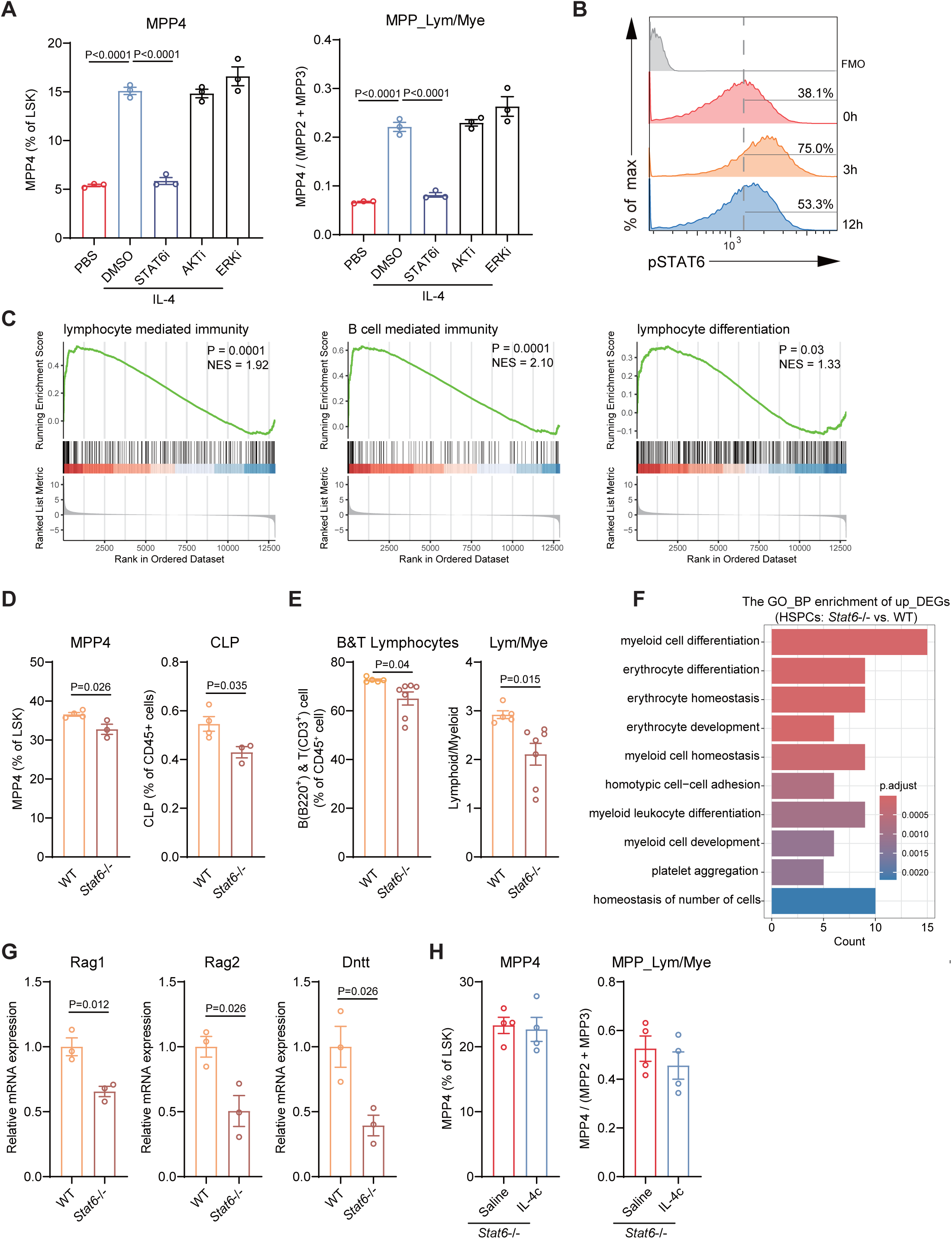
The IL-4-STAT6 signaling axis regulates genes important for lymphopoiesis. **A**, Percentage of MPP4 within isolated HSPCs, and the MPP_Lym/Mye ratio, with or without pre-treatment using different inhibitors (STAT6i (AS1517499): 50 nM, AKTi (MK-2206): 50 nM, ERKi (PD98059): 20 µM) and 50 ng/ml IL-4 *in vitro* (n=3). **B**, Flow cytometry plots of the intercellular staining of pSTAT6 on HSPCs from mice treated with IL-4c for different time (FMO, fluorescence minus one). **C**, GSEA enrichment profiles of gene sets associated with different lymphoid-related pathways. **D**, Percentage of MPP4 within HSPCs, and percentage of CLP in bone marrow CD45^+^ cell from WT and *Stat6*^−/−^ mice (n=3-4). **E**, B and T lymphocytes and lymphoid-to-myeloid ratio in peripheral blood of WT and *Stat6*^−/−^ mice (n=5-7). **F**, Gene ontology (GO) pathway enrichment analyses of genes significantly upregulated in HSPCs from *Stat6*^−/−^ mice versus WT mice (log_2_FoldChange > 1, Padj<0.05). **G**, RT–qPCR analysis of *Rag1, Rag2* and *Dntt* gene expression in HSPCs of WT and *Stat6*^−/−^ mice (n=3). **H**, Percentage of BM MPP4 in HSPCs and the MPP_Lym/Mye ratio from *Stat6*^−/−^ mice with or without IL-4c injections (n=4). All data represent means ± s.e.m. Statistical significance was determined by one-way ANOVA with Tukey’s multiple-comparisons test (**A**) or unpaired two-tailed Student’s t-test (**D, E, G** and **H**).

STAT6 is an important transcription factor that regulates gene expression across various cell types^39,40^. Since the role of STAT6 in transcriptional regulation in HSPCs had not been previously characterized, we performed RNA sequencing (RNA-seq) on HSPCs isolated from saline– or IL-4c-treated mice. We found IL-4c treatment resulted in the upregulation of 268 genes and the downregulation of 107 genes (**Figure S3A**, **Table S1**). Gene-set enrichment analysis (GSEA) revealed significant positive enrichment of gene sets associated with lymphocyte-mediated immunity, B cell-mediated immunity, and lymphocyte differentiation (**Figure 2C**). Interestingly, *Flt3* and *Ly6d*, which encoding markers for MPP4 and B-lymphoid progenitors respectively, were significantly upregulated by IL-4 treatment *in vitro* (**Figure S3B**). In addition, the percentage of Ly6D^+^ cells in HSPCs were markedly increased, and this increase is dependent on STAT6 activity (**Figure S3C**).

To further validate the role of STAT6 in regulating lymphoid progenitor signature genes and lymphocyte differentiation, we next analyzed the hematopoietic lineage cells of *Stat6*-deficient (*Stat6*^−/−^) mice. We found that *Stat6*^−/−^ mice exhibited an increased percentage of LT-HSCs but no significant change in ST-HSC when compared to that of the control mice (**Figure S3D**). Analysis of progenitor populations revealed that STAT6 deficiency resulted in a significant reduction in lymphoid lineage cells, including MPP4 and CLP (**Figure 2D**), while increasing some myeloid progenitors, such as CMP and GMP (**Figure S3E**). Consistently, *Stat6*^−/−^ mice exhibited a reduced percentage of lymphocytes and lymphocyte/myeloid cell ratio (**Figure 2E**). To investigate transcriptomic changes in HSPCs of *Stat6*^−/−^ mice, we performed RNA-seq, which identified 116 upregulated genes and 134 downregulated genes in *Stat6*^−/−^ HSPCs (**Figure S3F**, and **Table S2**). Gene Ontology (GO) biological process enrichment analyses revealed that STAT6-deficient HSPCs primarily upregulate genes involved in myeloid cell differentiation and erythrocyte differentiation (**Figure 2F**). Further analysis of potential STAT6 downstream targets showed a significant reduction in genes essential for lymphocyte development, including *Rag1*, *Rag2*, and *Dntt*^41,42^ (**Figure 2G**). Notably, *Rag2*-deficient (*Rag2*^−/−^) mice also exhibited reduced MPP4 levels compared to WT controls (**Figure S3G**). Importantly, IL-4 administration failed to induce MPP transitions in *Stat6*^−/−^ mice, further demonstrating an essential role of STAT6 in IL-4–mediated HSPC lineage reprogramming (**Figure 2H**).

Because STAT6 can also be activated by other cytokines, such as IL-13, we next examined whether IL-13 regulates MPP conversion. *In vitro* treatment with IL-13 increased MPP4 frequency and promoted the MPP_Mye-to-MPP_Lym transition, but only at high doses and with much weaker effect compared to that of IL-4, which was effective at much lower doses (**Figure S3H**). Consistently, *in vivo* IL-13 administration also induced MPP conversion, but much less potent than IL-4 (**Figure S3I**). To assess functional consequences, we performed transplantation assays using HSPCs pretreated with IL-13. IL-13 exposure enhanced lymphoid differentiation in recipient mice, but again much less robust compared to IL-4 (**Figure S3J**).

Collectively, these findings demonstrate that STAT6 mediates IL-4–driven lymphoid bias in HSPCs by activating lymphoid-specific transcriptional programs, at least in part through regulation of Rag1 and Rag2, which are essential for lymphopoiesis.

### IL-4 promotes myeloid-to-lymphoid transition by acting on MPPs, but not HSCs

HSPCs are a heterogeneous cell population consisted of LT-HSCs, ST-HSCs (MPP1), MPP2, MPP3, and MPP4. Although IL-4 can induce a shift from myeloid-biased MPPs to lymphoid-biased MPPs, the specific cell population it affects is not clear. The IL-4-induced shift could take place in four different possibilities: 1) an increase in MPP4 proliferation; 2) a block of MPP4 to CLP differentiation; 3) an increase in HSC to MPP4 differentiation; and 4) an increase in MPP2&3 to MPP4 trans-differentiation (**Figure 3A**). Although IL-4c injection significantly increased the cell cycle activity across all HSPC subsets *in vivo*, IL-4 treatment *in vitro* did not affect cell cycle activity or total HSPC numbers, despite it increased the MPP4 cell numbers (**Figure S4A-S4B**, and **Figure S2H**). This suggests that IL-4 indirectly promotes cell cycle activity *in vivo*, and that proliferation is not the primary driver of the increased MPP4 percentage (the first possibility). Additionally, since downstream CLPs, BLPs, and mature B cells in peripheral blood are all increased following IL-4 treatment (**Figure 1F** and **1K**), the second possibility can also be ruled out.

**Figure 3.**
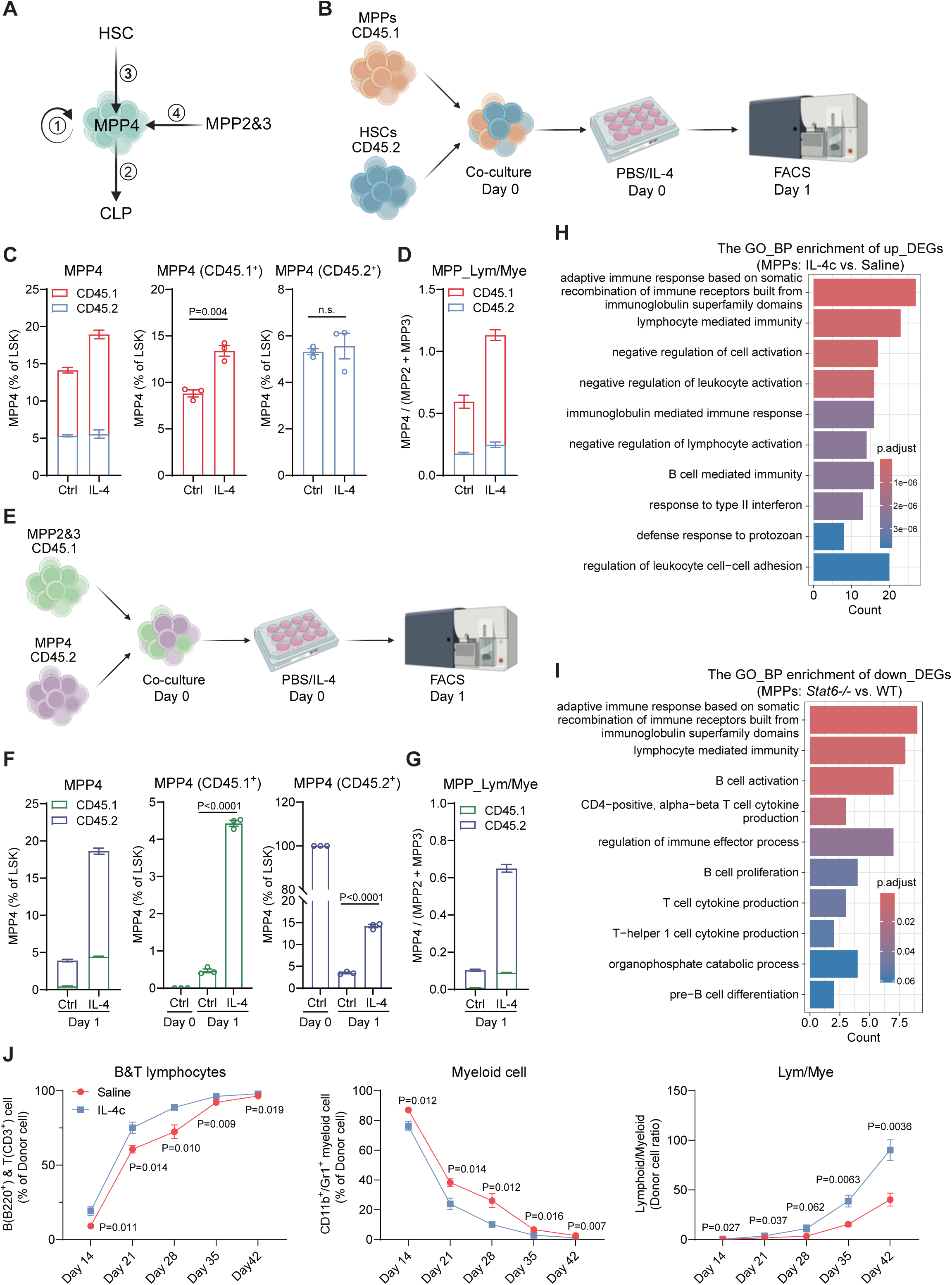
IL-4 promotes myeloid-to-lymphoid transition by acting on MPPs, but not HSCs. **A**, Diagram showing the different ways that could contribute to the change of MPP4 cell percentage/numbers. **B**, Experimental design for *in vitro* co-culture and treatment of CD45.1 MPPs and CD45.2 HSC. **C-D**, Percentage of MPP4 within co-cultured CD45.1 MPPs and CD45.2 HSC (**C**), and the MPP_Lym/Mye ratio (**D**), with or without treatment using 50 ng/mL IL-4 for 24 hours *in vitro*(n=3). **E**, Experimental design for *in vitro* co-culture and treatment of CD45.1 MPP2&3 and CD45.2 MPP4. **F-G**, Percentage of MPP4 within co-cultured CD45.1 MPP2&3 and CD45.2 MPP4 (**F**), and the MPP_Lym/Mye ratio (**G**), with or without treatment using 50 ng/mL IL-4 for 24 hours *in vitro* (n=3). **H**, GO pathway enrichment analyses of genes significantly upregulated in MPPs from IL-4c-treated versus saline-treated mice (log_2_FoldChange > 1, Padj<0.05). **I**, GO pathway enrichment analyses of genes significantly downregulated in MPPs from *Stat6*^−/−^ mice versus WT mice (log_2_FoldChange < –1, Padj<0.05). **J**, Transplantation of MPPs from IL-4c treated or untreated mice to lethally irradiated recipients: Donor-derived lymphoid cells (**J**, **left**), myeloid cells (**J**, **middle**) and lymphoid-to-myeloid ratio (**J**, **right**) in peripheral blood at the indicated days post-transplantation (n=5). All data represent means ± s.e.m. Statistical significance was determined by unpaired two-tailed Student’s t-test (**C-D**, **F-G** and **J**).

To determine whether the increase in MPP4 was due to HSC differentiation or MPP2&3 to MPP4 trans-differentiation, we utilized an *in vitro* culture system that allows direct tracing of different cell types (**Figure 3B**). To this end, we purified HSCs (LT-HSCs and ST-HSCs) from CD45.2 mice and MPPs (MPP2, MPP3, and MPP4) from CD45.1 mice and then mixed them together. After treating the mixed cells with IL-4 for 24 hours (**Figure 3B**), we analyzed their composition and observed an increase in both the percentage of MPP4 and the MPP_Lym/MPP_Mye ratio compared to that of the control group (**Figure 3C-3D**). Interesting, we found that only MPP4 from CD45.1 (MPP-derived cells), but not MPP4 from CD45.2 (HSC-derived cells) was increased after IL-4 treatment (**Figure 3C**). The increase in the MPP_Lym/Mye ratio was also predominantly contributed by CD45.1 cells (MPP-derived cells) (**Figure 3D**). To rule out potential differences in IL-4 responsiveness between the CD45.1 and CD45.2 mice derived cells, we conducted a similar experiment by mixing HSCs from CD45.1 mice and MPPs from CD45.2 mice, resulting in the same conclusion (**Figure S4C-S4D**). Taken together, these results confirm that the increased MPP_Lym/MPP_Mye ratio is not due to increased differentiation from upstream HSCs (the third possibility).

Having excluded three possible mechanisms, we concluded that trans-differentiation from MPP2&3 to MPP4 is likely the explanation. To gain support for this possibility, we isolated MPP4 and MPP2&3 from CD45.2 and CD45.1 mice, respectively, and mixed them together for IL-4 treatment (**Figure 3E**). The results showed that the increase in both MPP4 percentage and MPP_Lym/MPP_Mye ratio was contributed by both CD45.1 (MPP2&3-derived cells) and CD45.2 (MPP4-derived cells) cells (**Figure 3F-3G**). Isolated MPP4 rapidly lost their cell identity in culture medium, leading to a decrease in MPP4 cells after 24 hours (from 100% to ∼4%), but IL-4 treatment significantly mitigated this loss (∼15%, **Figure 3F, right panel**). Mixing MPP4 from CD45.1 mice and MPP2&3 from CD45.2 mice yield comparable results (**Figure S4E-S4F**). To gain further support *in vivo*, we analyzed the transcriptomic changes in MPPs and LT-HSCs separately after IL-4c treatment *in vivo*. RNA-seq analysis identified 289 up-regulated and 116 down-regulated genes in MPPs (**Figure S5A** and **Table S3**), and 235 up-regulated and 78 down-regulated genes in LT-HSCs, respectively (**Figure S5B**, and **Table S4**). GO and GSEA analysis of the differentially expressed genes (DEGs) showed that IL-4c upregulated genes are enriched for lymphocyte-related pathways in MPPs (**Figure 3H** and **Figure S5A**), while downregulated genes are enriched for myeloid leukocyte differentiation in MPPs (**Figure S5C**). However, the upregulated genes in IL-4c-treated LT-HSCs did not show significant enrichment in pathways related to lineage differentiation (**Figure S5B and S5D**). Additionally, RNA-seq analysis of MPPs from *Stat6*^−/−^ mice revealed 64 upregulated and 74 downregulated genes. In contrast to IL-4c-treated MPPs, the down-regulated genes in STAT6-deficient MPPs were enriched for lymphoid-related pathways (**Figure 3I** and **Figure S5E**, **Table S5**), while up-regulated genes are enriched for myeloid and platelet-related pathways (**Figure S5F**).

To further assess whether IL-4 treatment influences peripheral blood composition via affecting MPPs or LT-HSCs, we transplanted IL-4c-treated MPPs or LT-HSCs into recipient mice. Strikingly, transplantation of IL-4c-treated MPPs recapitulated the enhanced lymphoid contribution observed in whole HSPC transplants, mirroring the peripheral blood reconstitution patterns seen in earlier experiments (**Figure 3J**, compared to **Figure 1K**). In contrast, IL-4c-treated LT-HSCs exhibited no significant differences in lineage output compared to saline-treated controls (**Figure S5G**). Together, these findings support that IL-4 drives a myeloid-to-lymphoid shift in MPP differentiation by directly modulating their function, rather than acting through LT-HSCs. This MPP-specific effect underscores their pivotal role in IL-4-mediated lineage reprogramming.

### FLT3 potentiates IL-4–driven STAT6 activation for increased lymphoid commitment in MPPs

Next, we sought to investigate the molecular basis underlying the differential IL-4 signaling responses between MPPs and LT-HSCs. To this end, we conducted integrative analyses of published RNA-seq datasets^43^ comprising mouse LT-HSCs and different MPP subsets. Quantitative analysis revealed IL-4 downstream signaling components were broadly expressed across all HSPC subtypes with comparable levels (**Figure S6A**), indicating the involvement of additional regulatory elements modulating IL-4 signaling output between these populations. Comparative analysis of differentially expressed genes revealed a drastic upregulation of *Flt3* in MPPs (especially in MPP4) relative to its level in LT-HSCs (**Figure S6A**). Given that FLT3 is a class III receptor tyrosine kinase and that FLT3 ligand (FLT3L) deficiency has been associated with B-lymphocyte depletion in both mice and humans^44,45^, we hypothesized that FLT3 may enhance STAT6 phosphorylation and activation specifically in MPPs. To test this notion, we isolated HSPCs and exposed them to IL-4 stimulation *in vitro*. Flow cytometric analysis revealed significant increase in STAT6 Y641 phosphorylation in FLT3⁺ HSPCs compared to that in FLT3⁻ counterparts (**Figure 4A**). To gain further support for FLT3 mediated STAT6 activation, we co-expressed murine FLT3 and STAT6 in 293T cells. We found that expression of the full-length FLT3 enhanced STAT6 phosphorylation, and this effect depends on FLT3’s kinase domain and the effect is reduced by a FLT3 kinase inhibitor (**Figure 4B**). These results indicate that FLT3 kinase activity contributes to the IL-4–STAT6 signaling by enhancing STAT6 Y641 phosphorylation. In addition to STAT6, we examined other IL-4 downstream components, including JAK1 and IL-4Rα. We found that FLT3 phosphorylated IL-4Rα but not JAK1 (**Figure 4C**). Importantly, co-immunoprecipitation (co-IP) demonstrate that while no interaction between FLT3 and STAT6 was detected (data not shown), a strong association between FLT3 and IL-4Rα, the key receptor subunit that provides the docking site for STAT6 during canonical IL-4 signaling, was detected (**Figure 4D**). We next asked whether FLT3 activation alone could induce STAT6 phosphorylation. To this end, we treated primary HSPCs with FLT3L which led to robust STAT6 Y641 phosphorylation, confirming that FLT3 signaling is sufficient to activate STAT6 (**Figure 4E**). To assess whether our findings can be extended to humans, we first examined the effect of human FLT3 on IL-4R and STAT6 phosphorylation. By co-expressing human FLT3 with human IL-4R and STAT6, we found that FLT3 enhanced both IL-4R and STAT6 phosphorylation (**Figure 4F**), consistent with our findings in mice. The constitutively active FLT3 mutant (FLT3-ITD), commonly observed in AML (Acute Myeloid Leukemia) patients, also increased phosphorylation of IL-4R and STAT6 (**Figure 4F**). To validate these findings in hematopoietic cells, we compared two AML cell line: HL-60 (FLT3 wild-type) and MOLM-13 (FLT3-ITD mutant). Upon human IL-4 stimulation, MOLM-13 cells displayed markedly higher STAT6 phosphorylation compared to HL-60 cells, and this effect was largely abolished by treatment with a FLT3 inhibitor (**Figure 4G**). Collectively, these results demonstrate that FLT3 activity contributes to the activation of the IL-4R–STAT6 signaling in both mouse and human hematopoietic cells.

**Figure 4.**
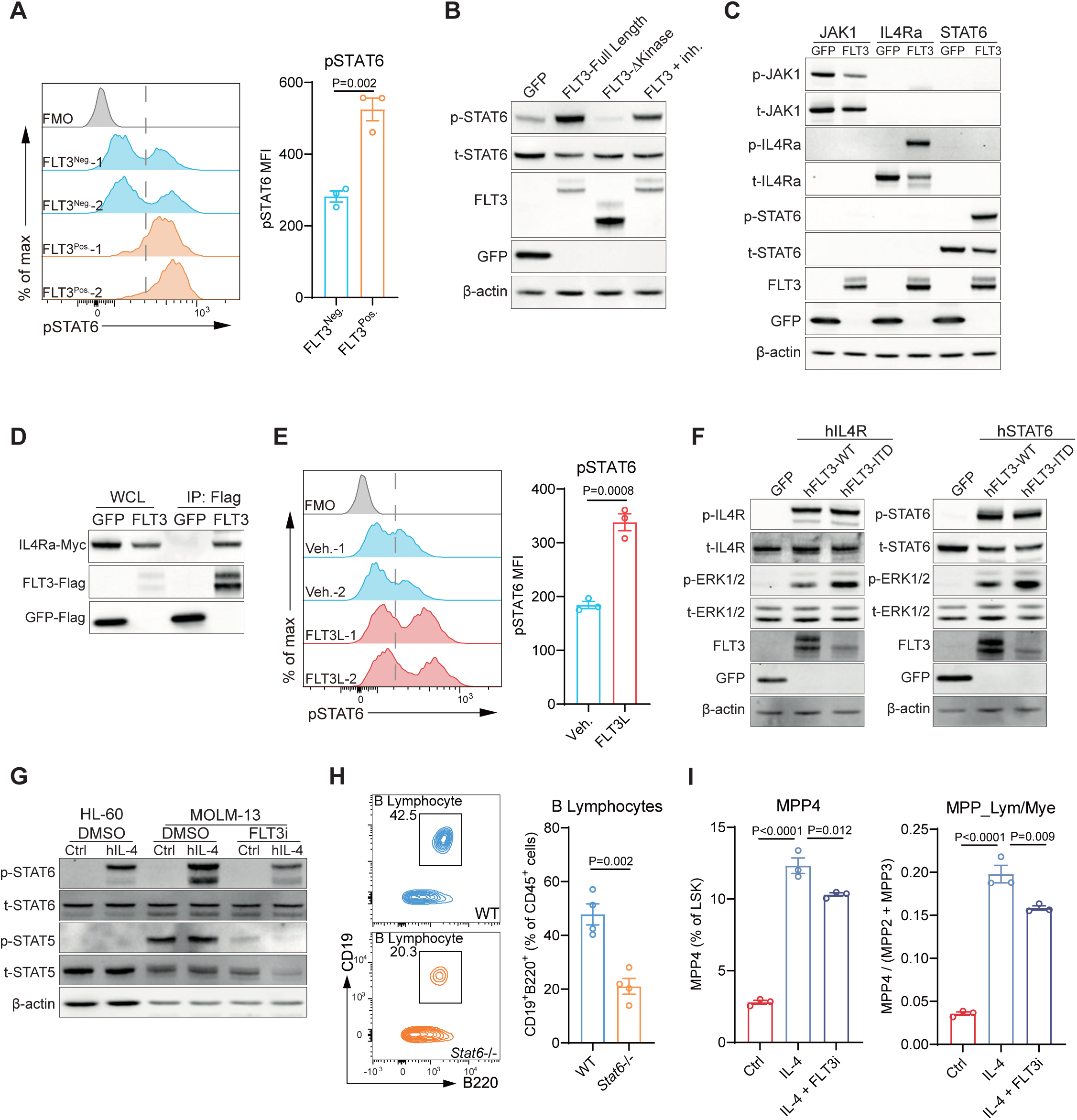
FLT3 potentiates IL-4–driven STAT6 activation for increased lymphoid commitment in MPPs. **A**, Representative FACS plots and quantification of the intercellular staining of pSTAT6 on FLT3^+^ and FLT3^−^ HSPCs treated with 50 ng/ml IL-4 *in vitro* for 30 minutes (n=3; FMO, fluorescence minus one; MFI, mean fluorescence intensity). **B**, Immunoblot analysis of 293T cells co-expressing *Stat6*-HA and *Il4ra*-Myc with *GFP*-Flag, *Flt3*(Full Length)-Flag, *Flt3*(ΔKinase)-Flag, or *Flt3*-Flag plus inhibitor (Quizartinib: 20nM) for 24 h. **C**, Immunoblot analysis of 293T cells co-expressing *JAK1*, *Il4ra* or *Stat6* with *GFP*-Flag or *Flt3*(Full Length)-Flag for 24 h. **D**, Immunoblot analysis of 293T cells co-expressing *Flt3*-Flag, *Stat6*-HA and *Il4ra*-Myc for 24 h and then co-IP were performed with anti-Flag antibody (IP, immunoprecipitation, WCL, whole-cell lysate). **E**, Representative FACS plots and quantification of the intercellular staining of pSTAT6 on HSPCs treated with 100 ng/ml FLT3L *in vitro* for 30 minutes (n=3). **F**, Immunoblot analysis of 293T cells co-expressing human *STAT6*-HA and *IL4R*-Myc with *GFP*-Flag, *FLT3*(WT)-Flag or *FLT3*(ITD)-Flag for 24 h. **G**, Immunoblot analysis of HL-60 and MOLM-13 cells treated with human IL-4 and/or FLT3 inhibitor (Quizartinib: 20 nM) for 1 hours. **H**, Representative FACS plots and quantification of the CD19^+^B220^+^ B lymphocytes differentiated from HSPCs at day 10 on OP9 stromal cells supplemented with FLT3L and IL-7 (n=4). **I**, Percentage of MPP4 within isolated HSPCs, and the MPP_Lym/Mye ratio, with or without pre-treatment using FLT3 inhibitor (Quizartinib: 20 nM) followed by 50 ng/ml IL-4 for 24 hours *in vitro* (n=3). All data represent means ± s.e.m. Statistical significance was determined by unpaired two-tailed Student’s t-test (**A**, **E, H** and **I**).

To functionally assess the relevance of the FLT3–STAT6 axis in lymphoid differentiation, we utilized the OP9 stromal co-culture system with FLT3L supplementation. We found that wild-type HSPCs efficiently generated CD19⁺B220⁺ lymphocytes, whereas the STAT6-deficient HSPCs exhibited a marked reduction in lymphoid output (**Figure 4H**). Finally, pharmacological FLT3 inhibition partially blocked IL-4–induced MPP transition (**Figure 4I**). Together, these findings demonstrate that FLT3 cooperates with IL-4 signaling to promote STAT6 phosphorylation and activation, thereby facilitating lymphoid commitment from MPPs.

### IL-4 suppresses inflammation-induced myelopoiesis by restoring the MPP_Lym/MPP_Mye ratio

Building on the mechanism identified above, which highlight the role of FLT3 in potentiating IL-4–driven STAT6 activation and lymphoid commitment under steady-state conditions, we next investigated whether IL-4 also modulates hematopoiesis during inflammatory stress. Given that IL-4 can specifically reverse IL-1β-induced myelopoiesis *in vitro* (**Figure 1A-1B**), we analyzed its effect on the expression of some key myelopoiesis regulators. We found that the expression of *Cebpd, Gata2*, *Csf3r*, and *Csf2ra* were significantly reversed following IL-4 treatment (**Figure 5A**). The pro-inflammatory cytokine IL-1β can reduce MPP4 numbers and inhibit lymphopoiesis, accompanied by a shift from MPP_Lym to MPP_Mye^23^. Such shift was largely reversed by the treatment with IL-4 (**Figure 5B**). To analyze the IL-4’s antagonistic effects on pro-inflammatory signals *in vivo*, we utilized an inflammation model by intraperitoneal lipopolysaccharide (LPS) injection. Similar to previous findings^23^, LPS treatment resulted in a decrease in MPP4 and the MPP_Lym to MPP_Mye ratio (**Figure 5C**). Importantly, this effect was largely reversed by co-administration of IL-4c (**Figure 5C**).

**Figure 5.**
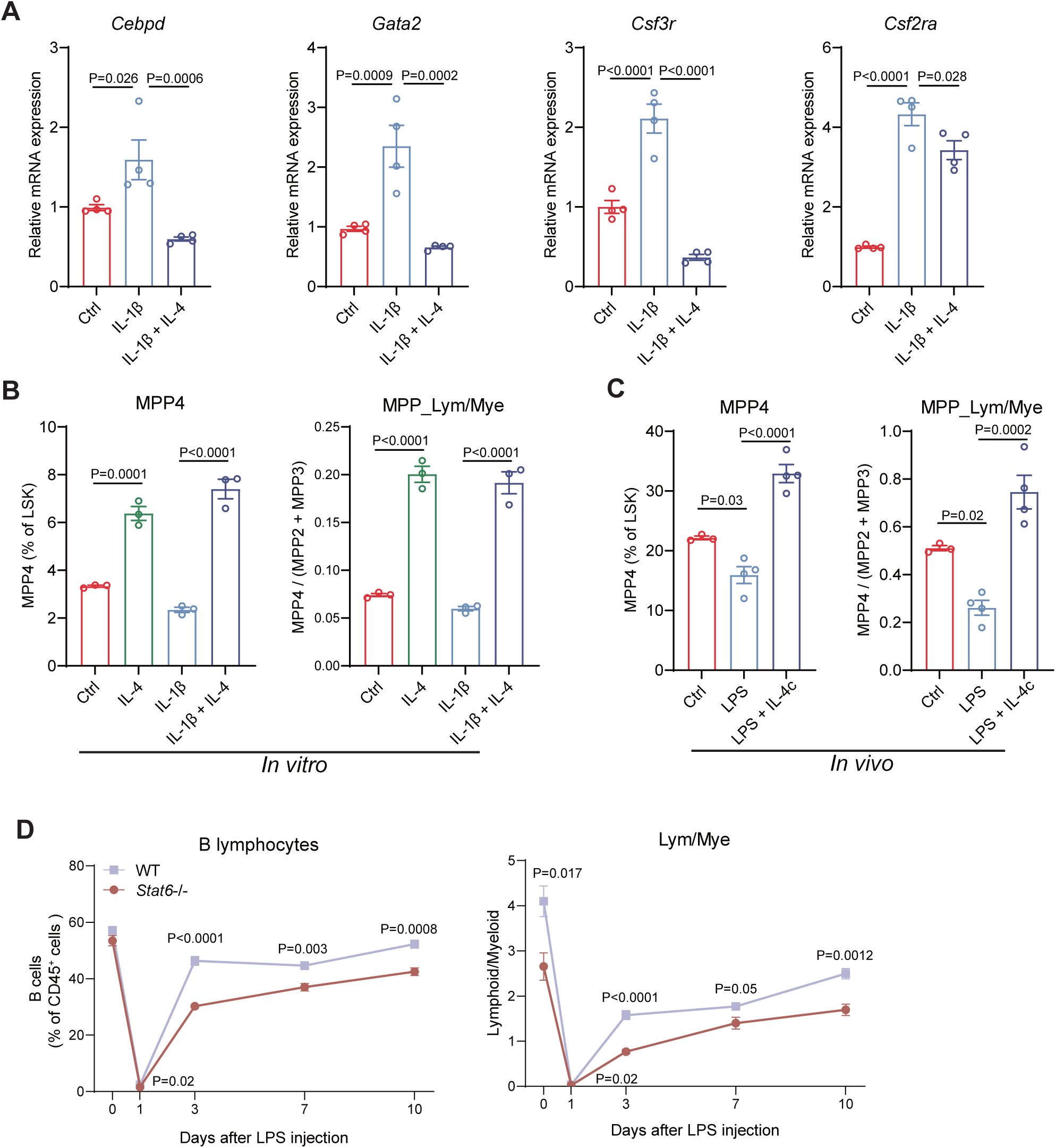
IL-4 suppresses inflammation-induced myelopoiesis by restoring the MPP_Lym/MPP_Mye ratio. **A**, RT–qPCR analysis of gene expression in myeloid differentiation assay with or without IL-1β/IL-4 treatment *in vitro* (n=4). **B**, Percentage of MPP4 within isolated HSPCs, and the MPP_Lym/Mye ratio, with or without 25 ng IL-1β pre-treatment for 24 hours followed by 50 ng/mL IL-4 for another 24 hours *in vitro* (n=3). **C**, Percentage of MPP4 within HSPCs, and the MPP_Lym/Mye ratio, from mice with or without treatment using 500 µg/kg LPS and/or IL-4c (n=3-4). **D**, B lymphoid cells and lymphoid-to-myeloid ratio in WT and *Stat6*^−/−^ mice after 500 µg/kg LPS injection for different days (n=5-6). All data represent means ± s.e.m. Statistical significance was determined by one-way ANOVA with Tukey’s multiple-comparisons test (**A-C**) or unpaired two-tailed Student’s t-test (**D**).

During the LPS-induced inflammatory response, not only pro-inflammatory cytokines but also some anti-inflammatory cytokines, including IL-4, were substantially upregulated^46^. To further evaluate IL-4’s role in regulating hematopoiesis during inflammation, we injected LPS into WT and *Stat6*^−/−^ mice and monitored changes in lymphoid and myeloid cells in peripheral blood. In both WT and *Stat6*^−/−^ mice, B-lymphoid cells decreased dramatically one day after LPS treatment (**Figure 5D**, **left panel**). However, while B-lymphoid cell recovery started by day three post-LPS treatment in WT mice, recovery in *Stat6*^−/−^mice was significantly delayed (**Figure 5D**, **left panel**). Consistently, lymphoid-to-myeloid ratio also exhibited a similar pattern (**Figure 5D**, **right panel**). These results demonstrate that IL-4 plays an important role in lymphopoiesis and lymphocyte recovery following inflammatory responses.

### IL-4 rejuvenates aged hematopoietic system by enhancing lymphopoiesis

Chronic inflammation is one of the defining hallmarks of aging, and is often regarded as a driver of myeloid-biased hematopoiesis^24,25,47,48^. This bias exacerbates inflammatory cytokine production, creating a positive feedback loop that perpetuates immune dysfunction and systemic decline. Such a shift in hematopoietic balance, favoring myeloid over lymphoid lineage differentiation, undermines the immune system’s ability to combat infections and maintain homeostasis^26–28^. Given that IL-4 can counteract inflammation-induced myeloid bias, we next examined whether IL-4 signaling contributes to aging-associated hematopoietic imbalances. To this end, we first measured IL-4 levels in bone marrow fluid of young (2–3 months) and aged (20 months) mice using ELISA and found no significant differences (**Figure S6B**). We then investigated downstream IL-4 signaling by integrative analysis of public RNA-seq datasets^43^ encompassing LT-HSCs, ST-HSCs, and MPP subsets of young and aged mice. We found that key genes of the IL-4 signaling pathway were significantly downregulated across various subsets of aged HSPCs, with the most pronounced reductions observed in MPPs (**Figure S6A**). To investigate IL-4 signaling in the context of human aging, we analyzed a recently published single-cell RNA-seq dataset of human HSPCs^49^. We found *FLT3*, *IL4R* and downstream STAT6 target gene (*RAG1* and *RAG2*) expression was markedly reduced in elderly HSPCs compared with adult HSPCs (**Figure S6C**), consistent with our findings in the mouse model. This downregulation likely contributes to the impaired hematopoietic function observed during aging.

To assess a potential restorative effect of IL-4 on aged HSPCs, we firstly administered a single dose of IL-4c to 18-month-old mice. One day after the administration, we analyzed the MPP composition and observed a substantial aging-associated decline in MPP4 subset as well as an overall skewing of MPP towards a myeloid bias, which were fully restored by IL-4c treatment to the levels observed in young mice, significantly reversed the MPP_Lym/Mye imbalance (**Figure 6A**). Consistently, IL-4c treatment also increased the MPP downstream lymphoid progenitors CLP and decreased the myeloid progenitor CMP, further supporting its potential in restoring lymphopoiesis in aged individuals (**Figure 6B**). To further investigate the molecular signature change of IL-4 treated aged MPPs, we performed RNA-seq on MPPs isolated from saline– or IL-4c-treated 18-month-old mice. IL-4c treatment resulted in the upregulation of 219 genes and downregulation of 84 genes (**Figure S6D**, **Table S6**). Consistently, comparative transcriptomic analysis of aged MPPs following IL-4c treatment revealed an upregulation of lymphoid-specific pathways and a downregulation of neutrophil-associated pathways, indicating a molecular shift toward lymphoid lineage commitment (**Figure 6C and Figure S6E**). In contrast, analysis of IL-4c treated aged LT-HSCs revealed upregulated cell cycle pathways but no significant changes in lineage differentiation pathways (**Figure S6F-S6G** and **Table S7**), which is similar to that observed in IL-4c-treated young LT-HSCs (**Figure S5B** and **S5D**). These results suggest that the aging-associated effect of IL-4c treatment is primarily mediated through its impact on MPPs rather than on HSCs.

**Figure 6.**
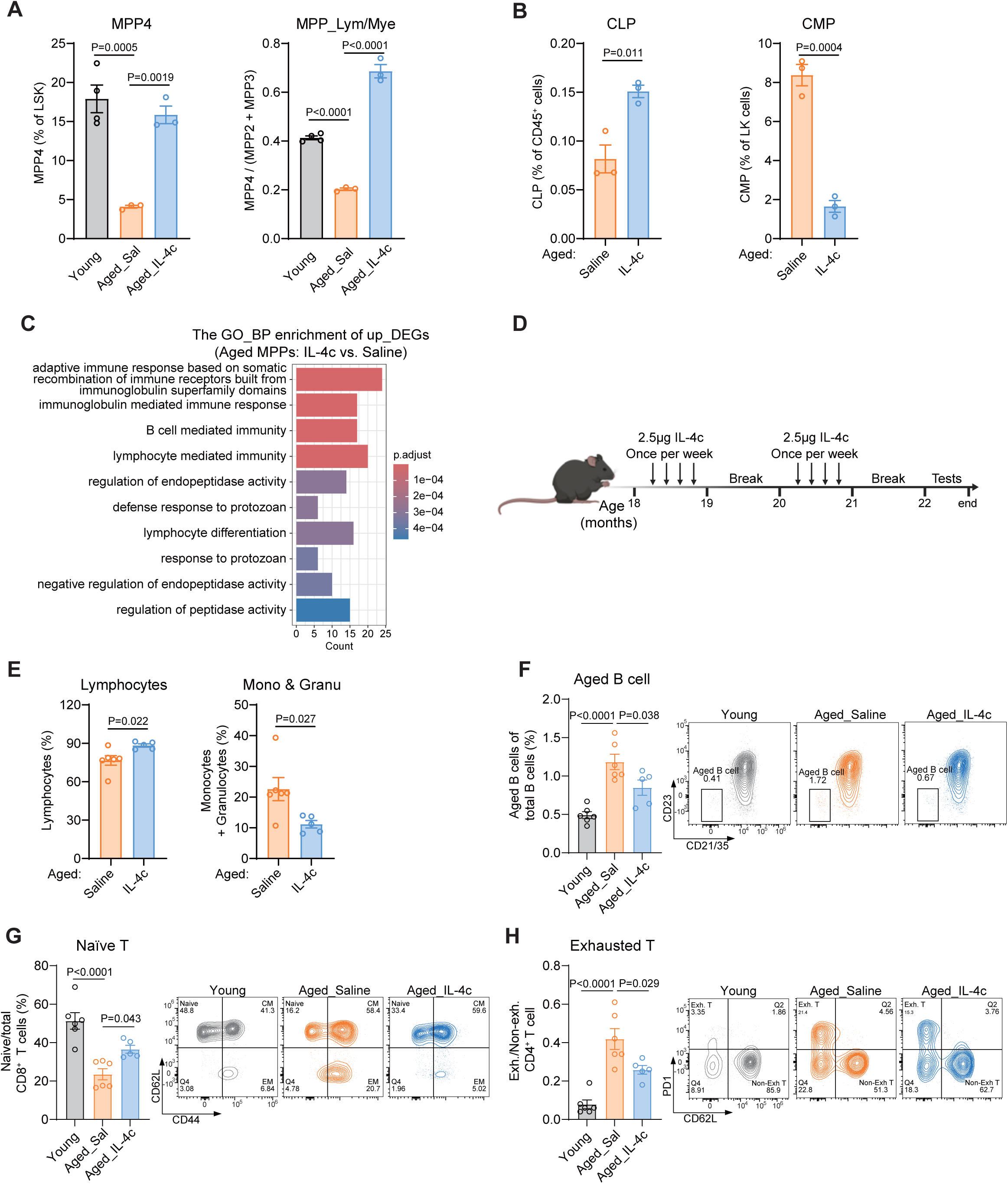
IL-4 rejuvenates aged hematopoietic system by enhancing lymphopoiesis. **A**, Percentage of MPP4 within HSPCs, and the MPP_Lym/Mye ratio, from young and aged mice with or without treatment using IL-4c (n=3-4). **B**, Percentage of CLP and CMP from aged mice with or without IL-4c injections (n=3). **C**, GO pathway enrichment analyses of genes significantly upregulated in MPPs from IL-4c-treated versus saline-treated aged mice (log_2_FoldChange > 1, Padj<0.05). **D**, Diagram showing the experimental design of long-term IL-4c treatment on aged mice. **E**, Complete blood counting analysis of lymphocytes, and monocyte & granulocytes (Mono & Granu) from aged mice with or without long-term IL-4c injections (n=5-6). **F-H**, Aged B cells (CD21/CD35^−^CD23^−^) as the percentage of total mature B cells (CD19^+^IgM^+^CD93^−^CD43^−^) in the blood (**F**); Naive CD8^+^ T cells (CD44^−^CD62L^+^) as the percentage of total CD8^+^ T cells in the blood (**G**); Exhausted CD4^+^ T cell ratio (percentage of PD1^+^CD62L^−^ cells)/(percentage of PD1^−^CD62L^+^ cells) (**H**) in the blood of aged mice with or without long-term IL-4c injections (n=5-6). All data represent means ± s.e.m. Statistical significance was determined by one-way ANOVA with Tukey’s multiple-comparisons test (**A** and **D-F**) or unpaired two-tailed Student’s t-test (**B** and **E**).

These encouraging results prompted us to assess the long-term effects of IL-4c using a multiple-dose regimen (**Figure 6D**). We first examined IL-4 dynamics following IL-4c injection and found that a single dose can maintain detectable serum IL-4 levels for more than 3 days (**Figure S6H**). Given that IL-4c–induced MPP transitions remained evident even 7 days after one injection (**Figure 1C–1E**), we choose administer IL-4c to aged mice once every 7 days. To minimize potential side effects, mice were given a one-month rest period between each month-long treatment cycle (four doses per cycle, for a total of eight doses). After an additional one-month rest to eliminate direct effects of IL-4c on mature peripheral blood cells, we performed whole-blood analysis. We found that the treatment increased lymphoid cells and decreased granulocytes and monocytes, without significantly affecting RBCs or platelets levels (**Figure 6E and Figure S6I**). Because IL-4 is known to drive type 2 inflammation, we also measured serum IgE levels following IL-4c treatment and found IgE levels were not affected, indicating that a one-month rest period is sufficient to prevent persistent IL-4–driven type 2 inflammatory responses (**Figure S6J**).

A major limitation of the aged immune system is the impaired generation and function of T and B lymphocytes^50^. Considering that IL-4c treatment in aged mice increased lymphocyte progenitors, we asked whether these changes were sufficient to influence B and T cell subtypes. We found that IL-4c treatment reversed aging-associated changes in immune cell subsets, restoring aged B cells (**Figure 6F**), naïve T cells (**Figure 6G**), and exhausted T cells (**Figure 6H**) to levels comparable to those in young mice. Collectively, these results demonstrate that IL-4 can rejuvenate the aged hematopoietic system by reversing myeloid-biased hematopoiesis, restoring lymphoid progenitor populations, and enhancing immune functions.

### IL-4 mitigates aging-associated decline in tissue functions

Hematopoietic system dysfunction is a critical contributor to whole-body aging, and targeting age-related changes in hematopoiesis counteracts and even reverses aging phenotypes^51–53^. As individuals age, shifts in HSPC differentiation often lead to imbalances in immune cell production, favoring myeloid over lymphoid lineages. This imbalance contributes to systemic inflammation, reduced immune competence, and impaired tissue regeneration^27^. By restoring the hematopoietic balance, IL-4 may have potential for alleviating age-related dysfunctions. Thus, we explored whether IL-4-mediated rebalancing of hematopoiesis in aged mice could translate into improvements in tissue and systemic functions. Muscle function decline, cognitive decline, and metabolic disorders are among the major hallmarks of aging that impact overall health and quality of life. To examine whether IL-4c treatment improves cognitive function, hippocampal-dependent learning and memory were assessed using the Y-maze test and novel object recognition (NOR) test following long-term intravenous administration of IL-4c (**Figure 6D**). The results indicate that IL-4c-treated aged mice have a significant preference for the novel arm and the novel object compared to the saline-treated controls (**Figure 7A-7B**), indicating an improved cognitive function.

**Figure 7.**
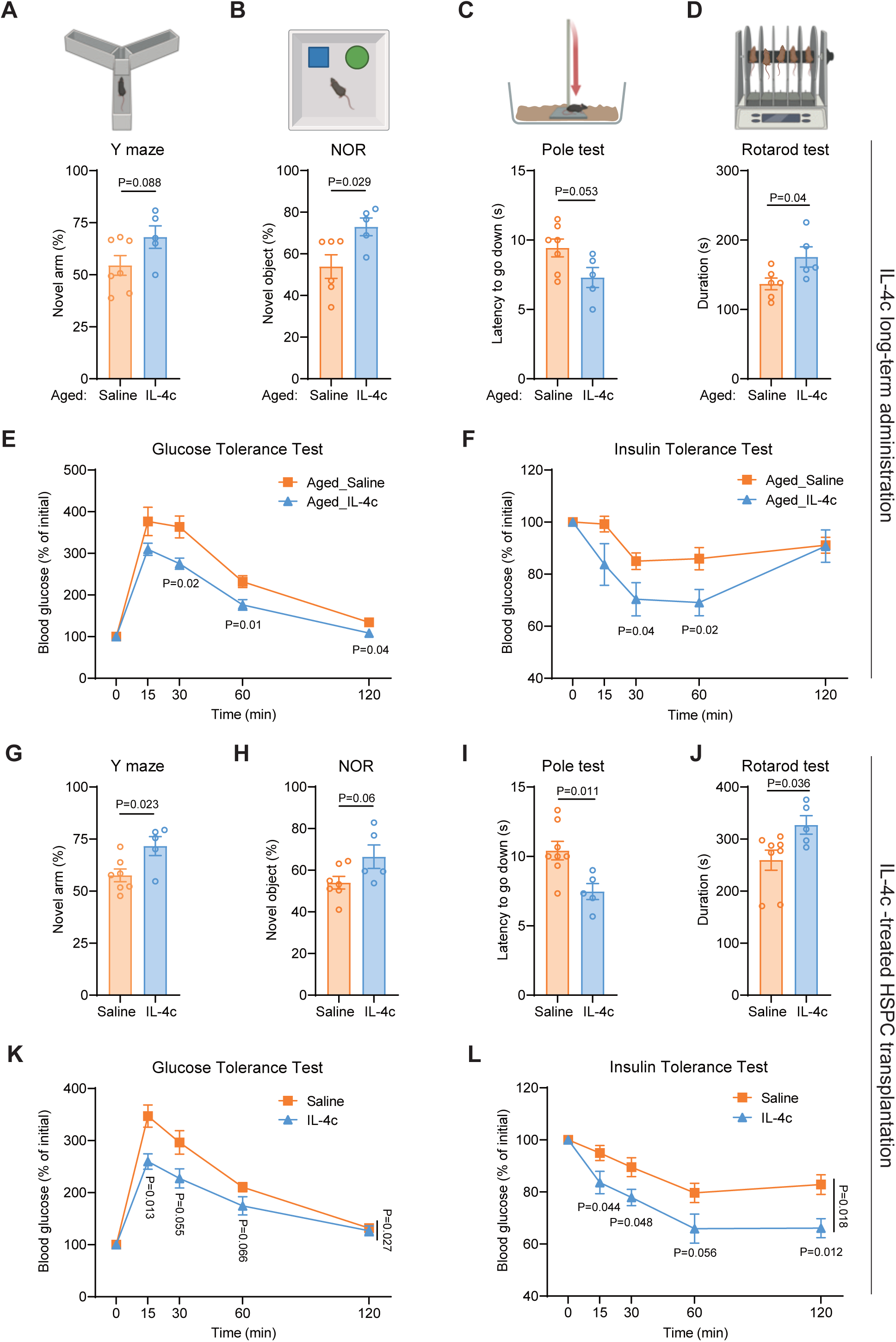
IL-4 treatment mitigates aging-associated decline in tissue function. **A-B**, Y maze (**A**) and novel object recognition (NOR, **B**) of long-term saline or IL-4c treated aged mice (n=5-7). **C-D**, Pole test (**C**) and rotarod test (**D**) of long-term saline or IL-4c treated aged mice (n=5-7). **E-F**, Insulin tolerance test (**E**) and glucose tolerance test (**F**) of saline or IL-4c treated aged mice (n=5-7). **G-J**, Y maze (**G**), NOR (**H**), Pole test (**I**) and rotarod test (**J**) of aged recipients transplanted with saline or IL-4c treated aged HSPCs (n=5-8). **K-L**, Insulin tolerance test (**K**) and glucose tolerance test (**L**) of aged recipients transplanted with saline or IL-4c treated aged HSPCs (n=5-8). All data represent means ± s.e.m. Statistical significance was determined by unpaired two-tailed Student’s t-test (**A-L**).

To assess the effects of IL-4c treatment on muscle function, we evaluated motor coordination, balance, and muscle strength using the pole test and rotarod test. We found that IL-4c-treated mice consistently outperformed saline-treated controls on both tests (**Figure 7C-7D**), highlighting significant improvements in motor function. Beyond enhancement of muscle and cognitive functions, metabolic dysfunctions, including glucose intolerance and type 2 diabetes, are prevalent in aging and are critical targets for intervention. Glucose tolerance tests (GTT) and insulin tolerance tests (ITT) revealed that IL-4c treatment significantly mitigated glucose intolerance and insulin resistance in aged mice (**Figure 7E-7F**). These findings suggest that IL-4 administration not only influences hematopoietic balance but also has systemic metabolic benefits. Given that inflammatory cytokines contribute to adipose tissue metabolic dysfunction^54^, our histological analyses showed that IL-4c–treated mice had markedly smaller adipocytes in both brown and subcutaneous white adipose tissue (**Figure S7A**), consistent with improved adipose remodeling. Furthermore, decreased lipid droplet accumulation was observed in the liver and bone marrow, suggesting enhanced fatty acid metabolism and reduced ectopic fat deposition (**Figure S7A**). Collectively, these histological and functional improvements highlight the potential of IL-4c as a therapeutic agent to combat aging-associated metabolic dysfunctions and functional decline.

Since IL-4 can directly influence peripheral immune cells as well as non-hematopoietic cell types, which may confound the interpretation of its tissue rejuvenation effects^55^, we performed a transplantation experiment to determine whether these benefits were hematopoietic in origin. To this end, aged HSPCs treated with either saline or IL-4c were transplanted into aged recipient mice, and one-month post-transplantation peripheral blood output was analyzed which revealed that IL-4c-treated HSPCs produced significantly more lymphoid cells and fewer myeloid cells compared to the saline controls (**Figure S7B, left and middle panel**). Additionally, increased chimerism rates in recipient mice transplanted with IL-4c-treated HSPCs were observed, indicating enhanced transplantation efficiency (**Figure S7B**, **right panel)**. Six months post-transplantation, mice receiving IL-4c-treated HSPCs exhibited improved cognitive and muscle function, mirroring the effects observed with direct IL-4c administration (**Figure 7G-7J**). Furthermore, glucose homeostasis was also improved in aged mice received IL-4c-teated HSPC transplantation (**Figure 7K-7L**). These results support the conclusion that the observed systemic benefits are predominantly driven by the restoration of HSPC lymphopoiesis, positioning IL-4c as a promising therapeutic agent in counteracting aging-associated metabolic and functional decline.

## Discussion

In this study, we provided compelling evidence that IL-4 functions as a potent modulator of hematopoiesis by counteracting pro-inflammatory effects. The capacity of IL-4 to influence hematopoietic progenitor cell fate under conditions of inflammation-induced dysregulation is particularly noteworthy. Through both *in vitro* and *in vivo* analyses, we showed that IL-4 shifts MPP from a myeloid-biased to a lymphoid-biased state via STAT6-dependent transcriptional reprogramming. This shift favors lymphopoiesis over myelopoiesis, effectively rebalancing hematopoietic lineage allocation. Mechanistically, this process is driven by the cooperative interaction between FLT3 and the IL-4 signaling pathway, which together enhance STAT6 activation and promote lymphoid differentiation from MPPs. These molecular changes extend beyond progenitor cells, leading to systemic improvements in immune function. Increased lymphoid cell output and enhanced adaptive immune potential in peripheral blood underscore IL-4’s ability to strengthen immune resilience in antagonizing cytokine-driven inflammation.

The ability of IL-4 to antagonize pro-inflammatory myelopoiesis holds significant implications for conditions characterized by chronic inflammation. Inflammatory conditions, such as autoimmune diseases and chronic infections, often lead to immune dysfunction characterized by skewed hematopoiesis and impaired immune responses^56,57^. By reversing IL-1β-mediated suppression of MPP4 and restoring lymphoid progenitor populations, IL-4 provides a counterregulatory mechanism to maintain hematopoietic equilibrium during immune challenges. Importantly, our LPS-induced inflammation model revealed that endogenous IL-4 contributes to lymphoid recovery post-inflammation, as neutralizing IL-4 delayed B lymphocyte repopulation. These findings align with emerging evidence that type 2 cytokines temper excessive inflammatory responses, but extend this paradigm by demonstrating direct effects on HSPC fate decisions. Moreover, the systemic benefits of IL-4-mediated modulation of hematopoiesis may also enhance the efficacy of other therapeutic strategies, including immunotherapies, by optimizing the immune cell repertoire. Additionally, IL-4’s therapeutic potential extends to mitigating hematopoietic dysregulation associated with aging. Chronic low-grade inflammation, or “inflammaging,” is a hallmark of the aging process and is known to skew hematopoiesis toward a myeloid bias^47,58^. The transcriptomic analyses of young and aged HSPCs revealed downregulation of IL-4 signaling components, particularly in MPP subsets, suggesting that age-related IL-4 deficiency exacerbates myeloid bias. Strikingly, IL-4 administration or IL-4-treated HPSC transplantation in aged mice not only restored MPP4 levels and lymphoid output but also improved cognitive function, motor coordination, and glucose metabolism. This positions IL-4 as a multifaceted therapeutic agent to combat both hematopoietic and functional decline in aging.

The cellular source of IL-4 in the bone marrow niche is critical for its role in regulating hematopoiesis. Several studies indicate that innate immune cells, particularly basophils and eosinophils, are major sources of IL-4. For example, the IL-4 eGFP expression in the bone marrow primarily originates from eosinophils and basophils^59^, and the bone marrow basophils have also been shown to secrete IL-4 in response to IL-33^60^. Similarly, mast cells, basophils, and eosinophils are programmed for IL-4 expression early in ontogeny^61^. Analysis of single-cell RNA-seq data^62,63^ confirmed that basophils are the predominant source of *Il4* mRNA in the bone marrow under steady-state conditions. Emerging evidence suggests that HSPCs themselves may also contribute to IL-4 production. RNA-seq data from our lab and an independent study^43^ indicate that HSCs and MPPs express *Il4*, with levels declining during aging, suggesting a role for HSPC-derived IL-4 in modulating the niche. Together, these findings support that IL-4 in the bone marrow is primarily from basophils, with potential contributions from HSPCs, and highlight its role in orchestrating hematopoietic balance and immune recovery.

While IL-4 plays a critical role in hematopoietic regulation, several important questions remain regarding its broader implications and potential applications. One is the long-term impact of IL-4 treatment on HSPC dynamics. Prolonged modulation of HSPCs *in vivo* raises the risk of stem cell exhaustion, which could compromise hematopoietic sustainability over time^64^. In addition, the direct effects of IL-4 on downstream progenitors—including CLPs, GMPs, MEPs, and CMPs—remain poorly understood. In tumor settings, IL-4 signaling in the bone marrow can promote immunosuppressive myelopoiesis through GMPs, thereby influencing tumor progression and outcomes^59^. Interestingly, although IL-4 treatment in mice increases GMP numbers, a similar effect is observed in STAT6-deficient mice, suggesting that IL-4’s influence on progenitors may involve STAT6-independent pathways or more complex regulatory mechanisms that need further investigation.

It is also important to contextualize these findings within the clinical landscape of IL-4–targeted therapies. IL-4R blockade (e.g., dupilumab) is widely used for treating atopic diseases without noticeable acceleration in hematopoietic aging. This apparent paradox likely reflects different biological contexts: atopic diseases are driven by pathological, systemic excess of type 2 cytokines (IL-4 and/or IL-13), whereas aging is characterized by a physiological deficiency of homeostatic IL-4 signaling within the HSPCs. Consequently, therapeutic blockade may normalize hyperactive signaling in peripheral tissues without fully abrogating the basal signaling required for HSPC function. Furthermore, the hematopoietic system exhibits significant cytokine redundancy. Other cytokines, such as IL-7, may exert some level of compensatory effects to sustain lymphopoiesis when IL-4 signaling is dampened. This compensatory potential likely preserves immune stability in patients undergoing IL-4 blockade.

### Limitation of the Study

Beyond hematopoiesis, IL-4 has been reported to promote tumor progression by inducing M2 macrophage polarization, driving tissue fibrosis, and fostering resistance to immune checkpoint blockade (e.g., anti–PD-1)^65–68^. These pro-tumorigenic activities highlight a major limitation for systemic IL-4–based therapies. While our findings suggest that IL-4 protects the aging hematopoietic system by counteracting myeloid bias, its broad immunomodulatory functions raise several concerns. First, systemic IL-4 delivery may not only accelerate tumor growth but also provoke type 2 inflammatory responses, worsening conditions such as asthma, allergy, or fibrosis. Second, the heterogeneity of IL-4R expression across tissues increases the risk of unintended activation in barrier or peripheral organs. To address these limitations, therapeutic strategies must prioritize targeted delivery. Approaches such as bone marrow–restricted IL-4 administration, nanoparticle-mediated delivery, or hematopoietic multipotent progenitor – specific IL-4R agonists could enhance hematopoietic benefits while minimizing off-target effects. Importantly, although our data show that reduced IL-4 responsiveness in HSPCs contributes to hematopoietic aging, the translational application of IL-4 restoration will require careful preclinical validation in tumor-prone and inflammation-prone settings. A context-specific and compartmentalized interventions might provide a favorable therapeutic window.

## Resource availability

### Lead contact

Requests for further information, resources, and reagents should be directed to and will be fulfilled by lead contact, Yi Zhang.

### Materials availability

Biological reagents generated for the purpose of this study can be requested from the lead contact.

### Data and code availability

The RNA-seq datasets of young and aged hematopoietic stem and progenitor cells are obtained from GSE162607^43^. Raw and processed RNA-seq data generated in this study are publicly available in the Gene Expression Omnibus (GEO) repository (accession numbers: GSE290821). Any additional information required to reanalyze the data reported in this paper is available from the lead contact upon request.

## Acknowledgements

We thank Dr. Ting Yan of Brigham and Women’s Hospital for analyzing the human HSPC scRNA-seq data, and the support of BCH FICR-PCMM Flow and Imaging Cytometry Resource and its director Dr. Viraga Haridas, as well as Mouse Behavior Core of Harvard Medical School and its director Dr. Barbara Caldarone. This project was supported by the Milky Way Research Foundation and the Howard Hughes Medical Institute (HHMI). Y.Z. is an investigator of the HHMI.

## Author contributions

Y.Z. and J.Y. conceived the project; J.Y. designed the experiments; J.Y. and Y.W. performed the transplantation; J.Y. performed most of the other experiments; J.Y. and Y.Z. discussed and interpreted the results, and wrote the manuscript.

## STAR methods

### EXPERIMENTAL MODEL AND PARTICIPANT DETAILS

#### Mice

All experiments were conducted in accordance with the National Institute of Health Guide for Care and Use of Laboratory Animals and approved by the Institutional Animal Care and Use Committee (IACUC) of Boston Children’s Hospital and Harvard Medical School. Mice were housed in a controlled environment with a temperature of 22 ± 1℃, humidity of 60% ± 10%, and a 12-hour light-dark cycle. For IL-4c injections, male C57BL/6 mice aged 2-3 months (Jackson Lab #00664) or 18 months were used. For HSPC or MPPs transplantation, donor mice were 2-3-month-old male CD45.2 C57BL/6 mice (Jackson Lab #00664), recipient mice were male B6 CD45.1 mice (Jackson Lab #002014), and helper cells were also sourced from male B6 CD45.1 mice. For experiments involving aged HSPC transplantation, donor mice were 24-month-old GFP^+^ male mice (Jackson Lab #006567), recipient mice were 18-month-old B6 CD45.2 male mice and helper cells were sourced from age-matched B6 CD45.2 mice. *Stat6*^−/−^ mice (C57BL/6, Jackson Lab #005977) and *Rag2*^−/−^ mice (C57BL/6, Jackson Lab #008449) were obtained from The Jackson Laboratories.

#### Physical Function Tests

In the pole test, mice were positioned at the top of a 50 cm grooved metal pole (1 cm diameter) with their heads pointing downward. The time taken to descend to the base (forelimb contact with the platform) was recorded using a stopwatch. This test was repeated twice after a 30-minute rest period, and the average descent time was calculated for each mouse. For rotarod test, mice were placed on an accelerating rotarod (Ugo Basile Apparatus) starting at 4 rpm. On the first day, mice were trained at 4 rpm for 5 minutes in two separate sessions. On the test day, the rotarod accelerated from 4 rpm to 40 rpm over 5 minutes. The time and speed at which the mice fell or completed two passive rotations were recorded. Each mouse completed three tests over two days, and the results were averaged. For Y-Maze test, mice were pre-acclimated in a separate holding room for 30 minutes. The procedure is consisted of a 3-minute habituation phase with one arm blocked, followed by a 3-minute test phase after a 2-minute intertrial interval. The maze was cleaned and the obstruction removed before the test phase and mice were placed into the start arm for the test trial. Distance and time traveled were recorded using Noldus EthoVision XT17 software. The percentage of time spent with the novel arm was calculated as: [(time with novel arm) / (time with trained arm + time with novel arm)] × 100%. For the novel object recognition test, mice explored an empty arena for 10 minutes (open field testing). After a 30-minute break, two identical objects were placed in the arena, and the mice were allowed to explore for 10 minutes. Following another 30-minute break, one object was replaced with a novel object, and mice explored for 10 minutes. The time spent exploring each object was quantified using Noldus EthoVision XT17. The percentage of time spent with the novel object was calculated as: [(time with novel object) / (time with trained object + time with novel object)] × 100%.

#### In vivo assays

For most IL-4c treatment, young or aged mice were injected intravenously once with either saline or 2.5 µg IL-4 (PeproTech #214-14) complexed with 12.5 µg of anti-IL-4 (Bio X Cell, #BE0045) for 24 hours. For long-term IL-4c treatment, 18-month-old mice were injected intravenously with IL-4c once a week for 4 weeks. After 4 weeks of treatment break, the mice were treated for another 4 weeks and another 4-week breaks, then peripheral blood, as well as physical and cognitive functional tests were performed. For IL-13 treatment, mice were injected intravenously once with either saline or 2.5 µg IL-13 for 24 hours (PeproTech # 210-13). For LPS-induced inflammation experiments, WT and *Stat6^−/−^*mice were injected intraperitoneally with 0.5 mg/kg LPS (Sigma-Aldrich, #L2630) in saline, then the peripheral blood was analyzed at different days as indicated. For LPS and IL-4c co-treatment, mice were injected intraperitoneally with 0.5 mg/kg LPS at day 0, then 2.5 µg IL-4 was injected intravenously at day 4. The mice were sacrificed and HSPCs were analyzed at day 5. For young mice transplantation experiments, recipient mice (2-3-month-old, CD45.1) were lethally irradiated (9.5 Gy) and injected intravenously with 5,000 donor (2-3-month-old, CD45.2) HSPCs, MPPs or LT-HSCs together with CD45.1 BM cells as helper. For aged mice transplantation experiments, recipient mice (18-month-old, CD45.2) were lethally irradiated (4.75 Gy × 2) and intravenously injected with 10,000 HSPCs isolated from 24-month-old GFP^+^ CD45.2 donor mice together with CD45.2 BM cells as helper. Prior to transplantation, donor HSPCs were treated with saline or IL-4c weekly for 4 weeks.

#### Flow cytometry

Bone marrow cells were harvested by crushing tibias, femurs, and pelvic bones. Lineage-positive cells (CD4, CD8, Gr-1, CD11b, B220, and Ter119) were removed using selection beads (STEMCELL Technologies #19856) to enhance sorting efficiency. Cells were stained with specific antibodies for analysis. For HSCs or MPPs, anti-Lineage-BV421, anti-c-Kit-PE, anti-Sca-1-PerCP-Cy5.5, anti-CD135/FLT3-PE-Cy7, anti-CD48-FITC, anti-CD150-APC were used. For myeloid progenitors, anti-Lineage-BV421, anti-c-Kit-PE, anti-Sca-1-PerCP-Cy5.5, anti-CD150-APC, anti-CD34-FITC and anti-FcγR-APC-Cy7 and anti-CD41-PE-Cy7 were used. For identification of CLP, a separate staining including anti-Lineage-BV421, anti-c-Kit-PE, anti-Sca-1-PerCP-Cy5.5, anti-CD135/FLT3-PE-Cy7, anti-CD48-FITC, anti-CD150-APC and anti-IL7R-APC-Cy7 were used. For myeloid differentiation *in vitro*, anti-c-Kit-PE, anti-Sca-1-PerCP-Cy5.5, anti-CD16/32-APC-Cy7 and anti-CD11b-FITC were used. For cell cycle analysis, lineage-positive BM cells were first stained with anti-Lineage-BV421, anti-c-Kit-PE, anti-Sca-1-PerCP-Cy5.5, anti-CD135/FLT3-PE-Cy7, anti-CD48-FITC and anti-CD150-APC, and then fixed and permeabilized with Cytofix/Cytoperm buffer (BD Biosciences) for 20 min on ice. After washing with Perm/Wash (BD Biosciences), cells were stained with anti-Ki67-AF700 in Perm/Wash for 30 min on ice, washed with Perm/Wash and then re-suspended in Perm/Wash containing 1 mg/ml DAPI (BD Pharmingen, 564907) before analysis. Non-cycling cells were defined as G0: Ki67^Neg^DAPI^Low^, while all remaining cells in G1, S, G2, and M phases were classified as cycling cells. For transplantation experiments, PB chimerism and lineage distribution of donor-derived cells were assessed by staining with anti-CD45.1-BV421, anti-CD45.2-PE-Cy7 or anti-CD45.2-APC-Cy7, anti-B220-PerCP-Cy5.5, anti-CD3-APC, anti-Gr-1-PE and anti-CD11b-PE. For T cell subset analysis, cells were stained with anti-CD3-APC-Cy7, anti-CD8-FITC, anti-CD4-BV510, anti-CD62L-BV711, anti-CD44-BV605, anti-CD25-PerCP-Cy5.5, anti-Foxp3-eFluor 450 and anti-PD1-PE-CF594. For B cell subset analysis, cells were stained with anti-CD19-PE, anti-IgM-FITC, anti-CD43-APC, anti-CD93-PE-Cy7, anti-CD23-PerCP-Cy5.5, anti-CD21/CD35-APC-Cy7, anti-CD45R/B220-PE-CF594 and anti-IgD-eFluor 450. For lymphoid differentiation assays with OP9 stoma cells, the cocultured cells were analyzed with anti-CD45-PE-Cy7, anti-CD19-PE, anti-CD11b-FITC, anti-Gr-1-APC and anti-B220-PerCP-Cy5.5 antibodies. For Phospho-STAT6 staining, lineage-positive cells (CD4, CD8, Gr-1, CD11b, B220, and Ter119) were removed using selection beads (STEMCELL Technologies #19856). Then Lin-cells were stained with antibodies against surface markers, including anti-c-Kit-AF700, anti-Sca-1-AF647, anti-CD135/FLT3-BV421. After fixed with 2% PFA for 20 mins, the cells were permeabilized with ice-cold 90% methanol for 30 mins. Samples were washed three times in PBS, then the collected cell pellet was resuspended in 100 µl of diluted anti-Phospho-Stat6 (Tyr641)-FITC in staining buffer. For human lymphoid-primed multipotent progenitor (LMPP) analysis, cultured CD34⁺ HSPCs were stained with anti-CD34-FITC, anti-CD38-BV421, anti-CD45RA-APC, anti-CD90-PE, and anti-CD135/FLT3-PE-Cy7, with LMPPs defined as CD34⁺CD38⁻CD90⁻CD45RA⁺FLT3⁺. Flow cytometry analysis was performed on the FACSCanto (BD Biosciences) or Cytek Aurora (Cytek Biosciences). Cell sorting was performed using a SH800S Cell Sorter (Sony Biotechnology).

#### *In vitro* assays

All cultures were performed at 37℃ in a 5% CO_2_ incubator. For HSPC culture, sorted LSK (Lin-Sca-1+cKit+) cells were plated in 96-well plates with ∼5,000 cells/well in 100 μL volume of cell culture media with Ham’s F-12 Nutrient Mix containing 1× PSG, 10 mM HEPES, 1 mg/ml PVA, 1 × ITSX, 100 ng/ml TPO (PeproTech), 10 ng/ml SCF (PeproTech). For IL-4 treatment to analyze MPP transition, cells were treated with 50 ng/mL IL-4 (or the indicated concentrations) for 24 h. For inhibitor assays, inhibitors (concentrations indicated in figure legends) were added 12 h before IL-4 treatment and maintained during the 24 h culture. For myeloid differentiation, 500-1000 LT-HSCs were sorted per well of a 96-well plate and cultured for 6-7 days. Cells were grown in StemSpan™ SFEM medium (STEMCELL Technologies, #09600) supplemented with 1 x penicillin/streptomycin and 2 mM L-glutamine, 25 ng/ml SCF, 25 ng/ml FLT3L, 25 ng/ml IL-11, 10 ng/ml IL-3, 10 ng/ml GM-CSF, 4 U/ml Epo and 25 ng/ml TPO (All from Peprotech). Cytokines were refreshed every two days by replacing 50% of the media. For pre-B colony formation assays, BM cells (2 x 10^5^ per 1.1 ml per 3 cm dish) were cultured in methylcellulose (STEMCELL Technologies, #M3630) and colonies were scored and counted after 7 days. For lymphoid differentiation assays with OP9 stoma cells, isolated HSPCs (3,000–4,000 cells/well in 24-well plates) were seeded into wells containing 80-100% confluent OP9 stromal cells. The co-cultured cells were grown in α-MEM medium supplemented with 20% FBS, 2 mM L-glutamine, penicillin/streptomycin, 5 ng/mL Flt3L and 5 ng/mL IL-7. The medium was half changed every 2 days. Differentiated lymphocytes were analyzed after 8-10 days culture by flow cytometry. For human CD34⁺ BM HSPC culture (STEMCELL Technologies, #70002.2), cells were thawed and expanded in StemSpan SFEM II supplemented with 1× StemSpan™ CD34+ expansion supplement and 500 nM UM729 for 2 days, followed by treatment with 50 ng/mL human IL-4 (Thermo Scientific, #200-04) for 24 h. Treated HSPCs were analyzed by flow cytometry.

#### Cell viability assay

HSPC viability was assessed by using Dead Cell Apoptosis Kits (Thermo Fisher Scientific, #V13241). HSPCs were washed once in cold PBS and resuspended in Annexin binding buffer. Samples were stained with Annexin V–AF488 and propidium iodide (PI) according to manufacturer’s instructions (final recommended dilution) for 10–15 min at room temperature in the dark. After the incubation period, 200 µL 1X annexin-binding buffer was added. Samples were analyzed on a flow cytometer and gated to exclude debris and doublets; populations were defined as live (Annexin V⁻ PI⁻), apoptotic (Annexin V⁺ PI⁻), and necrotic (Annexin V⁺ DAPI⁺).

#### Quantitative RT-PCR analysis

Total RNA was isolated from cells using Zymo Direct-zol RNA Miniprep kit. cDNA was synthesized with SuperScript III, and RT-PCR was performed with Fast SYBR Green Master Mix (Thermo Fisher) on ViiA 7 Real-Time PCR System (ThermoFisher). Relative expression level of mRNAs was calculated using the 2(-Delta Delta CT) method and *Actb* was used as an internal control.

#### Bulk RNA-seq library preparation

Bulk RNA-seq libraries were prepared using the Smart-seq2 method with the SMART-Seq v4 Ultra Low Input RNA Kit (Takara #634890) following the manufacturer’s instructions.

#### Histological analysis

Tissues (BAT, scWAT, liver and bone marrow) were fixed for 24 h in 4% paraformaldehyde (PFA) and subsequently stored in 70% ethanol. After the tissues were embedded in paraffin, the blocks were cut into 5-μm sections and stained with hematoxylin and eosin.

#### Immunoblot and immunoprecipitation

293T cells transfected with indicated plasmids for 24 hours were lysed in RIPA lysis buffer (Thermo Fisher), and protein extracts were resolved by NuPAGE 4–12% gel (Invitrogen, NP0322BOX) and transferred to NC membrane. Membranes blocked with protein blocking buffer were first incubated with the indicated primary antibodies, followed by horseradish peroxidase-conjugated anti-rabbit IgG or anti-mouse IgG secondary antibody. The signals were detected using an enhanced chemiluminescence (ECL) kit (Thermo Fisher Scientific) and imaged by iBright Imaging Systems (Invitrogen). Primary antibodies used included anti-HA (1:2,000, Cell Signaling Technology), anti-Myc (1:2,000, Bio-Rad), anti-Flag (1:2,000, Sigma-Aldrich), anti-STAT6 (1:2,000, Cell Signaling Technology), anti-Phospho-STAT6 (Tyr641) (1:2,000, Cell Signaling Technology), anti-IL-4R (1:2000, Cell Signaling Technology), anti-Phospho-IL-4R (Tyr497) (1:1000, Antibodies.com), anti-JAK1 (1:2000, Cell Signaling Technology), anti-Phospho-JAK1 (Tyr1034/1035) (1:1000, Cell Signaling Technology), and anti-β-actin (1:3,000, Cell Signaling Technology). Secondary antibodies used included goat anti-Rabbit IgG (H + L) superclonal secondary antibody-HRP (Thermo Scientific, 1:2,000) and goat anti-mouse IgG (H + L) secondary antibody-HRP (Thermo Fisher Scientific, 1:2,000). For immunoprecipitation experiments, cells transfected with the indicated plasmids were lysed in IP lysis buffer (Thermo Scientific) with cocktail protease inhibitors for 0.5 h at 4 °C. Cell lysates were then centrifuged for 10 min at 4 °C. The supernatant was incubated with Anti-Flag Magnetic Beads (MedChem Express) at 4 °C overnight. The immunoprecipitates were washed five times with immunoprecipitation lysis buffer and then analyzed by immunoblot.

#### Bone marrow IL-4 cytokine analysis

To collect bone marrow, both ends of the femur and tibia were cut to expose the marrow cavity. An 18 G needle was used to puncture the bottom of a 0.5 mL microcentrifuge tube, creating a small drainage hole. The bones were placed into the prepared tube with the knee end facing downward, and the tube was closed. This 0.5 mL tube was then nested inside a 1.5 mL microcentrifuge tube and centrifuged at 5,000 × g for 10 sec to release the marrow. The expelled marrow was resuspended in 100 µL PBS and centrifuged again at 3,00 × g for 3 min to remove cells and debris. The clarified supernatant was collected as bone marrow fluid, and IL-4 concentrations were measured using an ELISA kit according to the manufacturer’s protocol.

#### Glucose tolerance test and insulin tolerance test

For the GTT, mice were fasted for 12 h and then i.p. injected with D-(+)-glucose (Sigma-Aldrich) at a dose of 2 g per kg body weight. Blood glucose was measured from the tail bleeds at the indicated time after glucose injection. For the ITT, mice were fasted for 6 h and then i.p. injected with insulin (Eli Lilly) at a dose of 1 U per kg body weight. Blood glucose was measured from the tail bleeds at the indicated time after insulin injection.

#### RNA-seq data analysis

For bulk RNA-seq datasets, adaptor of all sequenced reads was first trimmed by Trim Galore. Then the filtered reads were mapped to the mouse genome (mm10) with HISAT2 (v2.0.4, https://daehwankimlab.github.io/hisat2/). Bam files were sorted and indexed in SAMtools (https://github.com/samtools/samtools/releases/). Assembly, quantification and normalization were performed in StringTie (v1.3.6, http://ccb.jhu.edu/software/stringtie/). DEseq2 (v1.18.1, https://bioconductor.org/packages/release/bioc/html/DESeq2.html) was used to analyze the DEGs. DEGs were subjected to GO (http://geneontology.org) and KEGG (https://www.genome.jp/kegg/) pathway enrichment analyses using the clusterProfiler R package (https://guangchuangyu.github.io/software/clusterProfiler/).

#### scRNA-seq analysis of human HSPCs

Single-cell RNA-seq data of human hematopoietic stem and progenitor cells (HSPCs) were obtained from GSE189161^49^. Data processing and analysis were performed in R using Seurat (v5.2) following the standard workflow^69^. To account for donor variability, datasets from different age groups were integrated using Seurat’s anchor-based integration approach. Cell lineage identities were assigned according to the annotations provided in the original study. Analyses of IL-4 signaling were carried out on the integrated dataset, focusing on HSPC lineage populations from adult (25–53 years old) and elderly (62–77 years old) donors. For visualization, bar plots comparing adult and elderly groups were generated in R (ggplot2). Log-normalized expression values for each gene were extracted at the single-cell level, and group means were plotted as bars with standard error. Statistical significance between groups was assessed using the Wilcoxon rank-sum test, with p-values displayed above the corresponding bars.

#### Statistical analysis

Sample sizes, replicates, and statistical tests are detailed in figure legends. Sample sizes were determined from prior or pilot experiments to ensure statistical validity while adhering to the 3R principles. Blinding was not done as all the analyses were performed quantitatively and not subjectively. Normal data distribution was assumed but not formally tested. Results are expressed as means ± s.e.m. and analyzed using Prism 8 (GraphPad) or Microsoft Excel 2019. Comparisons between two groups were performed with unpaired two-tailed Student’s t-tests, while one-way ANOVA with Tukey’s test was used for multiple group comparisons.

## List of Supplemental Tables

**Table S1.** DEGs of HSPCs isolated from mice treated with IL-4c versus saline. (Related to Figure 2)

**Table S2.** DEGs of HSPCs isolated from *Stat6*^−/−^ mice versus WT mice. (Related to Figure 2)

**Table S3.** DEGs of MPPs isolated from mice treated with IL-4c versus saline. (Related to Figure 3)

**Table S4.** DEGs of LT-HSCs isolated from mice treated with IL-4c versus saline. (Related to Figure 3)

**Table S5.** DEGs of MPPs isolated from *Stat6*^−/−^ mice versus WT mice. (Related to Figure 3)

**Table S6.** DEGs of MPPs isolated from aged mice treated with IL-4c versus saline. (Related to Figure 6)

**Table S7.** DEGs of LT-HSCs isolated from aged mice treated with IL-4c versus saline. (Related to Figure 6)

---

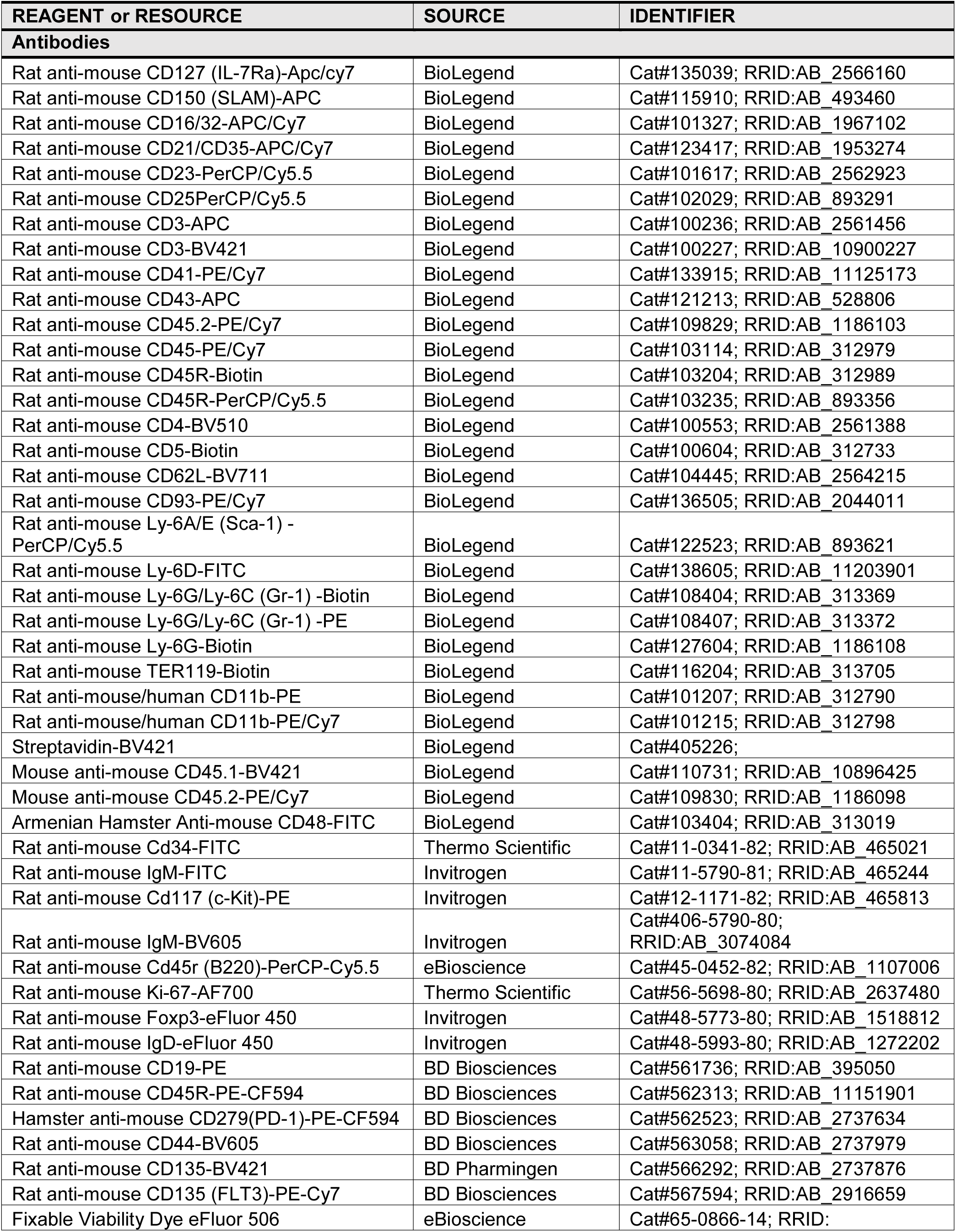
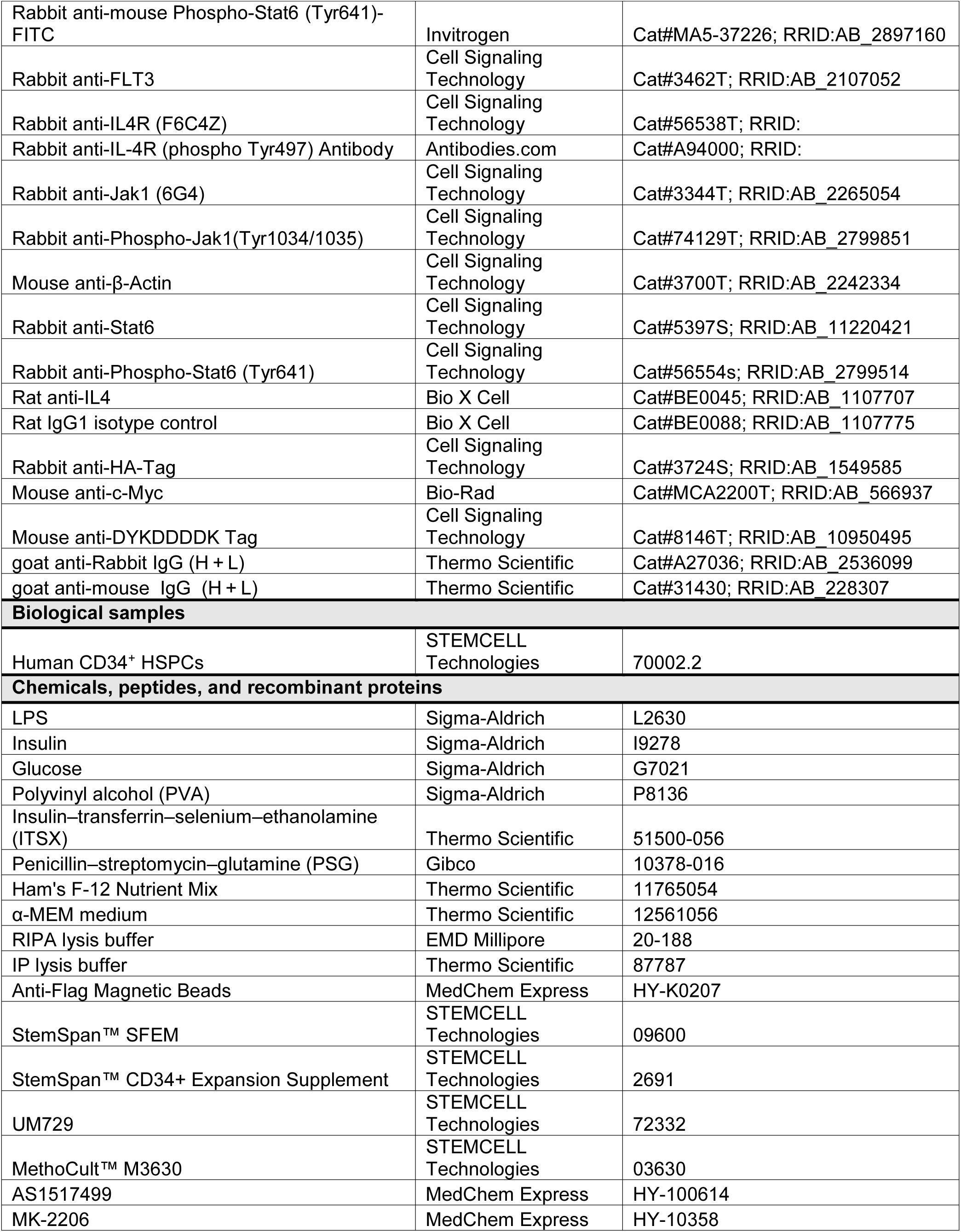
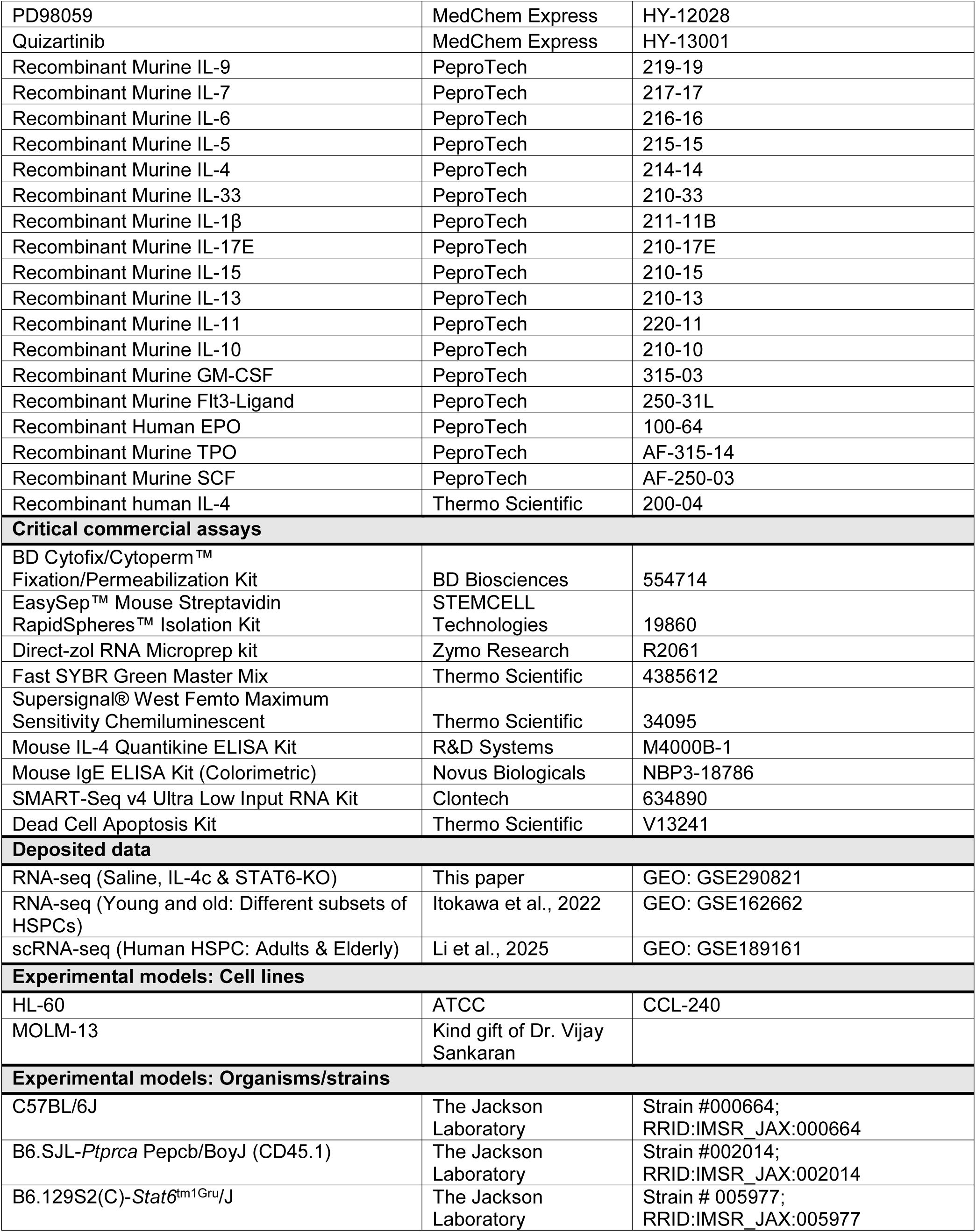
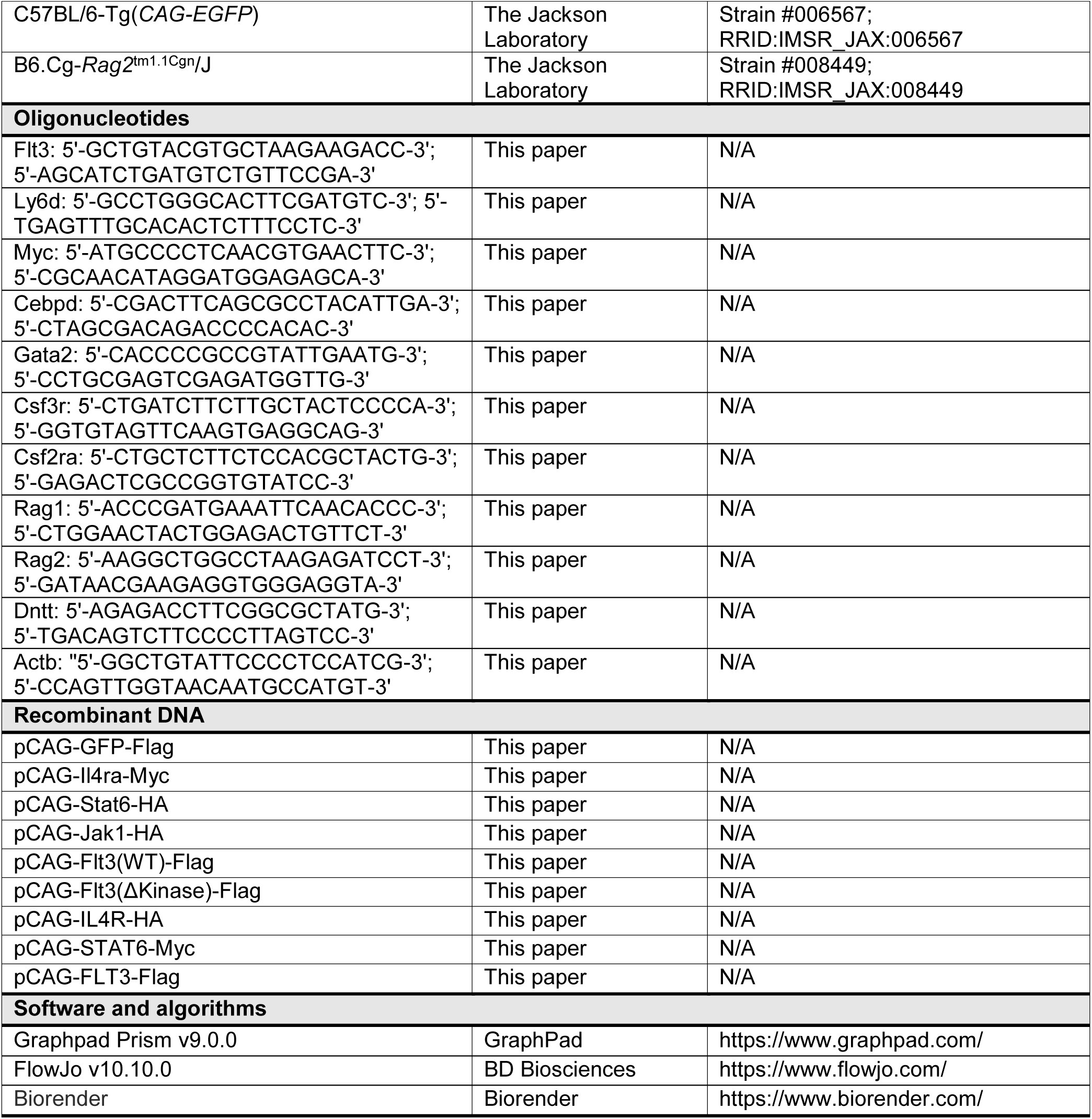
Key Resource Table.

## Supplemental Text and Figures

**Figure S1.**
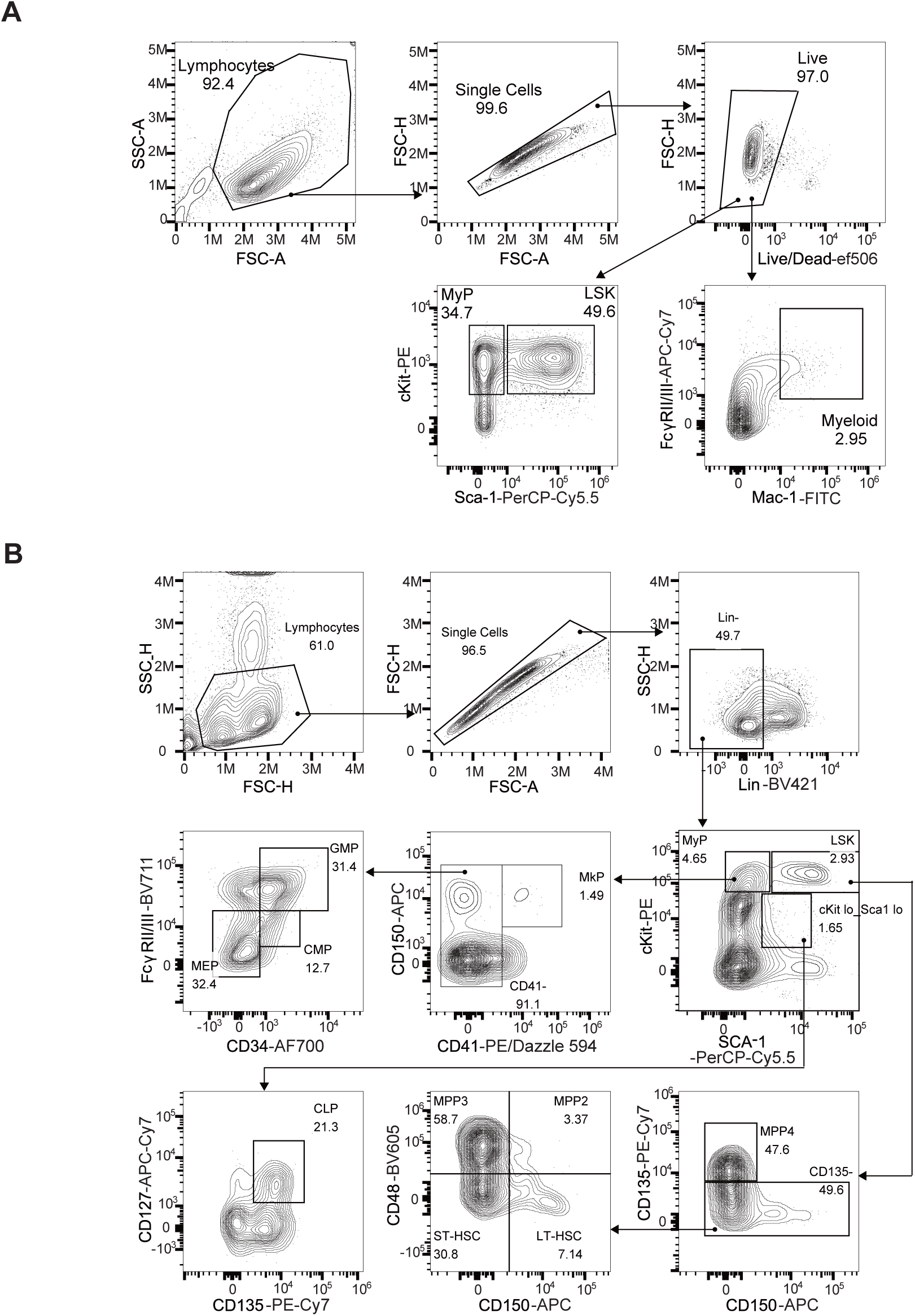
Gating strategy (Related to Figure 1) **A**, Representative gating strategy for *in vitro* myeloid cells differentiation assay. **B**, Representative gating strategy for BM progenitor populations.

**Figure S2.**
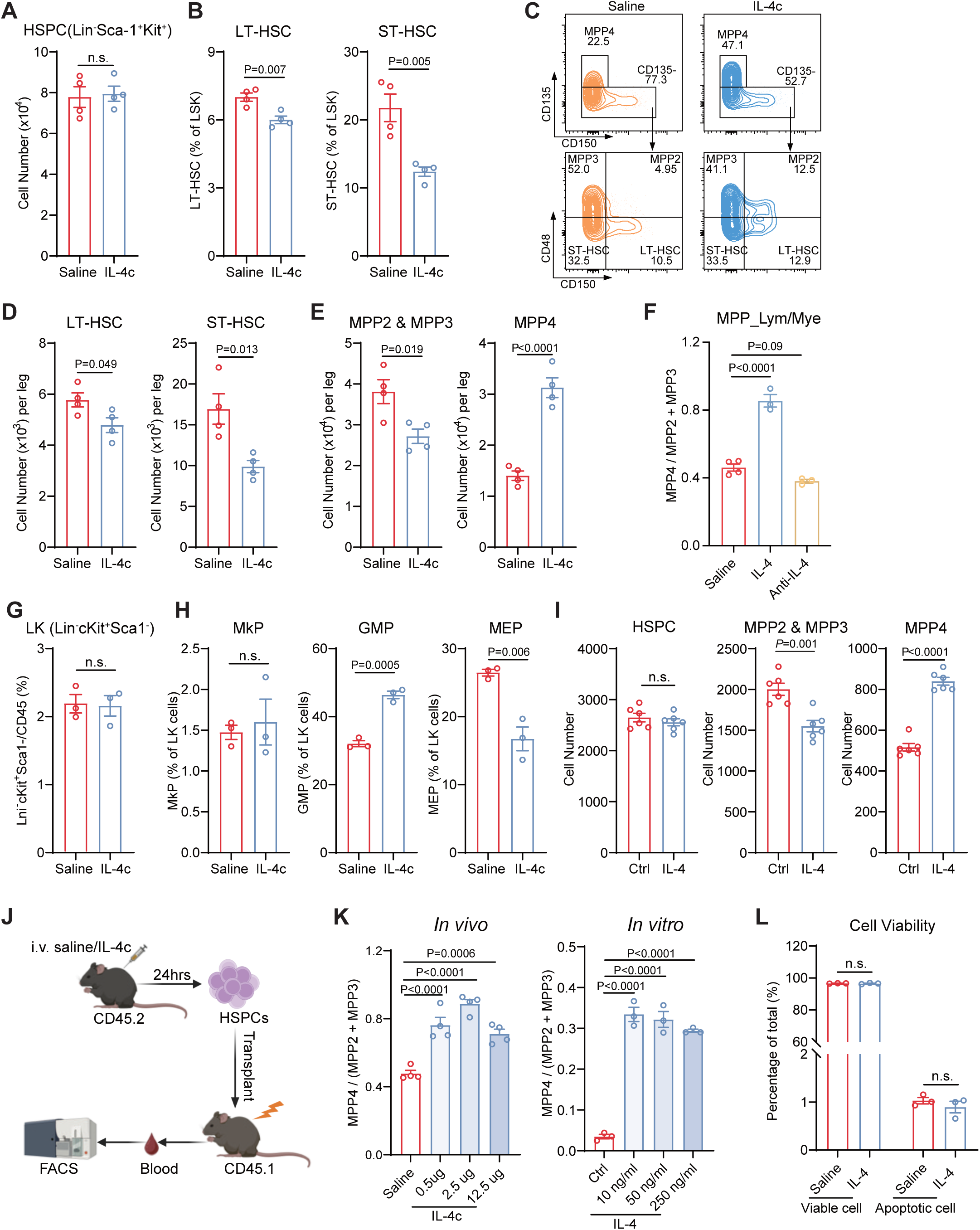
Effects of IL-4 treatment on hematopoietic stem and progenitor cells, and downstream cells (Related to Figure 1) **A**, Absolute numbers of the BM HSPCs from mice with or without IL-4c treatment (n=4). **B**, Percentage of BM LT-HSC and ST-HSC in HSPCs from mice with or without IL-4c treatment (n=4). **C**, Representative FACS plots of LT-HSC, ST-HSC, MPP2, MPP3 and MPP4 within HSPCs from mice with or without IL-4c treatment. **D-E**, Absolute numbers of LT-HSC and ST-HSC (**D**) as well as MPP2 & MPP3 and MPP4 (**E**) per leg with or without IL-4c treatment *in vivo* (n=4). **F**, The BM MPP_Lym/Mye ratio of WT mice treated with 2.5 ug IL-4 alone or 12.5 ug anti-IL-4 antibody alone for 24 hours (n=3-4). **G-H**, Percentage of BM LK populations (**G**), MkP, GMP and MEP (**H**) from mice with or without IL-4c treatment *in vivo* (n=3). **I**, Absolute numbers of the indicated populations (HPSCs, MPP2 & MPP3 and MPP4) with or without 50 ng/ml IL-4 treatment *in vitro* (n=6). **J**, Schematic of transplantation of HSPCs from IL-4c treated or untreated mice in lethally irradiated recipients. **K**, The MPP_Lym/Mye ratio after different doses of IL-4 treatment from *in vivo* (Left panel, n=4) and *in vitro* (Right panel, n=3). **L**, HSPC viability analysis after IL-4 treatment for 24 hours *in vitro* (n=3). All data represent means ± s.e.m. Statistical significance was determined by one-way ANOVA with Tukey’s multiple-comparisons test (**K**) or unpaired two-tailed Student’s t-test (**A-B**, **D-I** and **L**).

**Figure S3.**
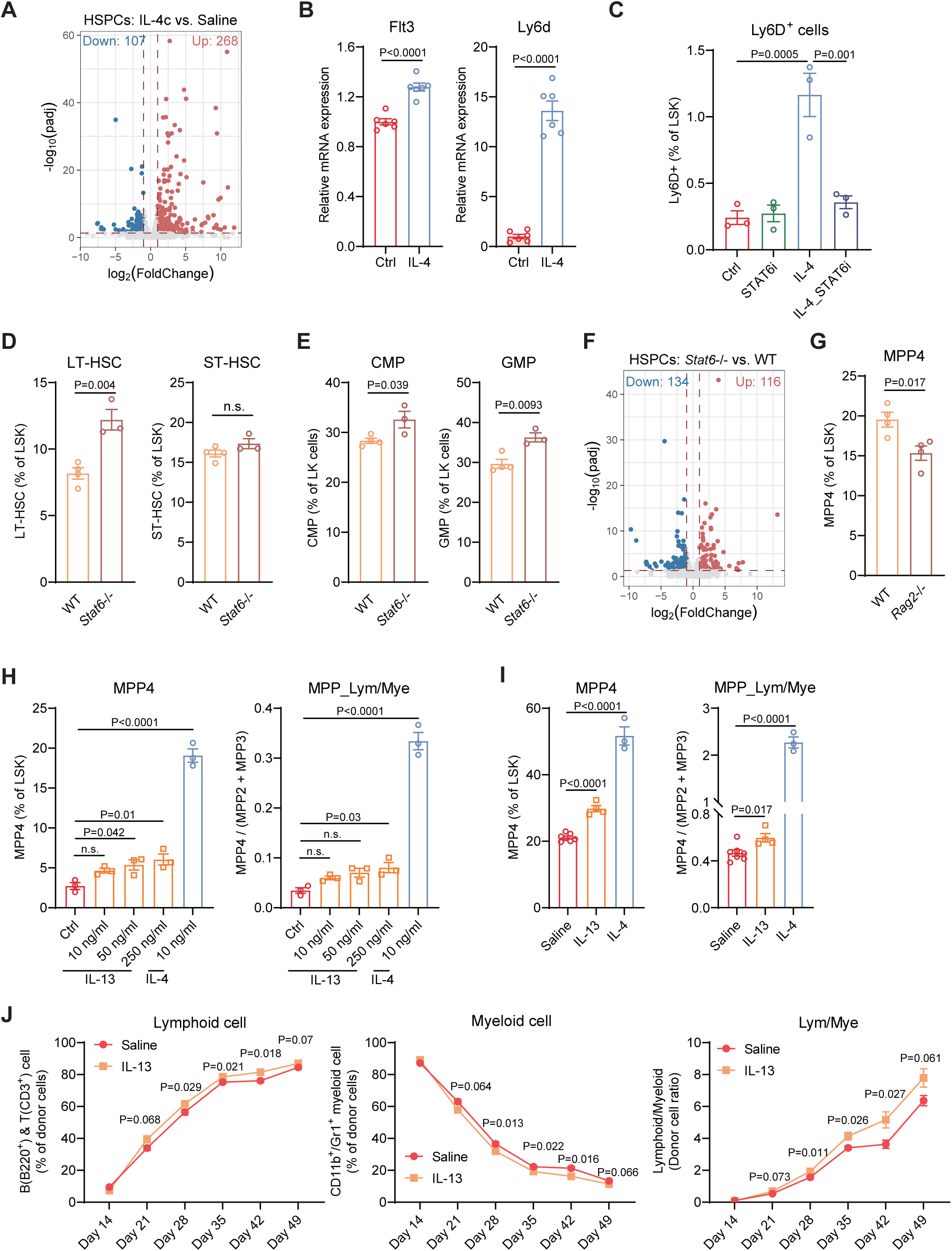
The STAT6 activation regulates genes important for lymphopoiesis (Related to Figure 2) **A**, Volcano plot showing the DEGs in HSPCs from IL-4c-treated versus saline-treated mice. **B**, RT–qPCR analysis of *Flt3* and *Ly6d* gene expression in isolated HSPCs treated with or without 50 ng/ml IL-4 for 24 hours *in vitro* (n=6). **C**, Percentage of Ly6D^+^ cells in isolated HSPCs with or without STAT6 inhibitor treatment *in vitro* (n=3). **D**, Percentage of BM LT-HSC and ST-HSC in HSPCs from WT and *Stat6*^−/−^ mice (n=3-4). **E**, Percentage of BM CMP and GMP in LK from WT and *Stat6*^−/−^ mice (n=3-4). **F**, Volcano plot showing the DEGs in HSPCs from *Stat6*^−/−^ versus WT mice. **G**, Percentage of BM MPP4 in HSPCs from WT and *Rag2*^−/−^ mice (n=4). **H**, Percentage of MPP4 subpopulations within isolated HSPCs and the MPP_Lym/Mye ratio, with or without 24 hours treatment using different concentration of IL-13 or IL-4 *in vitro* (n=3). **I**, Percentage of MPP4 subpopulations within HSPCs and the MPP_Lym/Mye ratio from WT mice with or without IL-13 or IL-4c injections (n=3-6). **J**, Transplantation of MPPs from IL-4c treated or untreated mice to lethally irradiated recipients: Donor-derived lymphoid cells, myeloid cells and lymphoid-to-myeloid ratio in peripheral blood at the indicated days post-transplantation (n=5). All data represent means ± s.e.m. Statistical significance was determined by one-way ANOVA with Tukey’s multiple-comparisons test (**C, H** and **I**) or unpaired two-tailed Student’s t-test (**B, D-E, G** and **J**).

**Figure S4.**
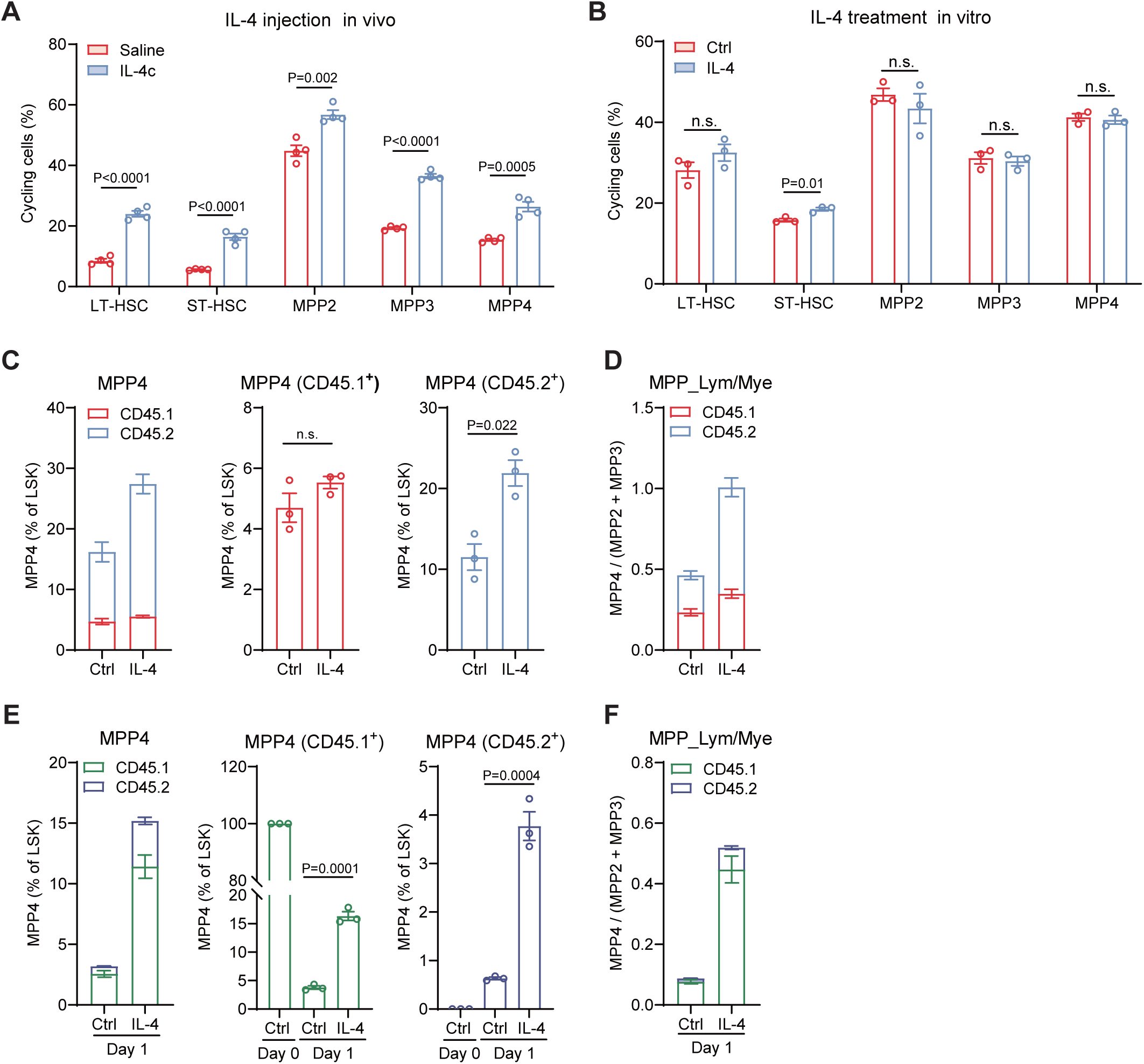
Effects of IL-4 treatment on HSPC proliferation and differentiation as well as MPP transition (Related to Figure 3) **A-B**, Percentage of cycling cells of indicated populations with or without IL-4c treatment *in vivo* (**A**) (n=4) or *in vitro* (**B**) (n=3). **C-D**, Percentage of MPP4 within co-cultured CD45.1 HSCs and CD45.2 MPPs (**C**), and the MPP_Lym/Mye ratio (**D**), with or without treatment using 50 ng/mL IL-4 for 24 hours *in vitro* (n=3). **E-F**, Percentage of MPP4 within co-cultured CD45.1 MPP4 and CD45.2 MPP2 & 3 (**E**), and the MPP_Lym/Mye ratio (**F**), with or without treatment using 50 ng/mL IL-4 for 24 hours *in vitro* (n=3). All data represent means ± s.e.m. Statistical significance was determined by unpaired two-tailed Student’s t-test (**A-F**).

**Figure S5.**
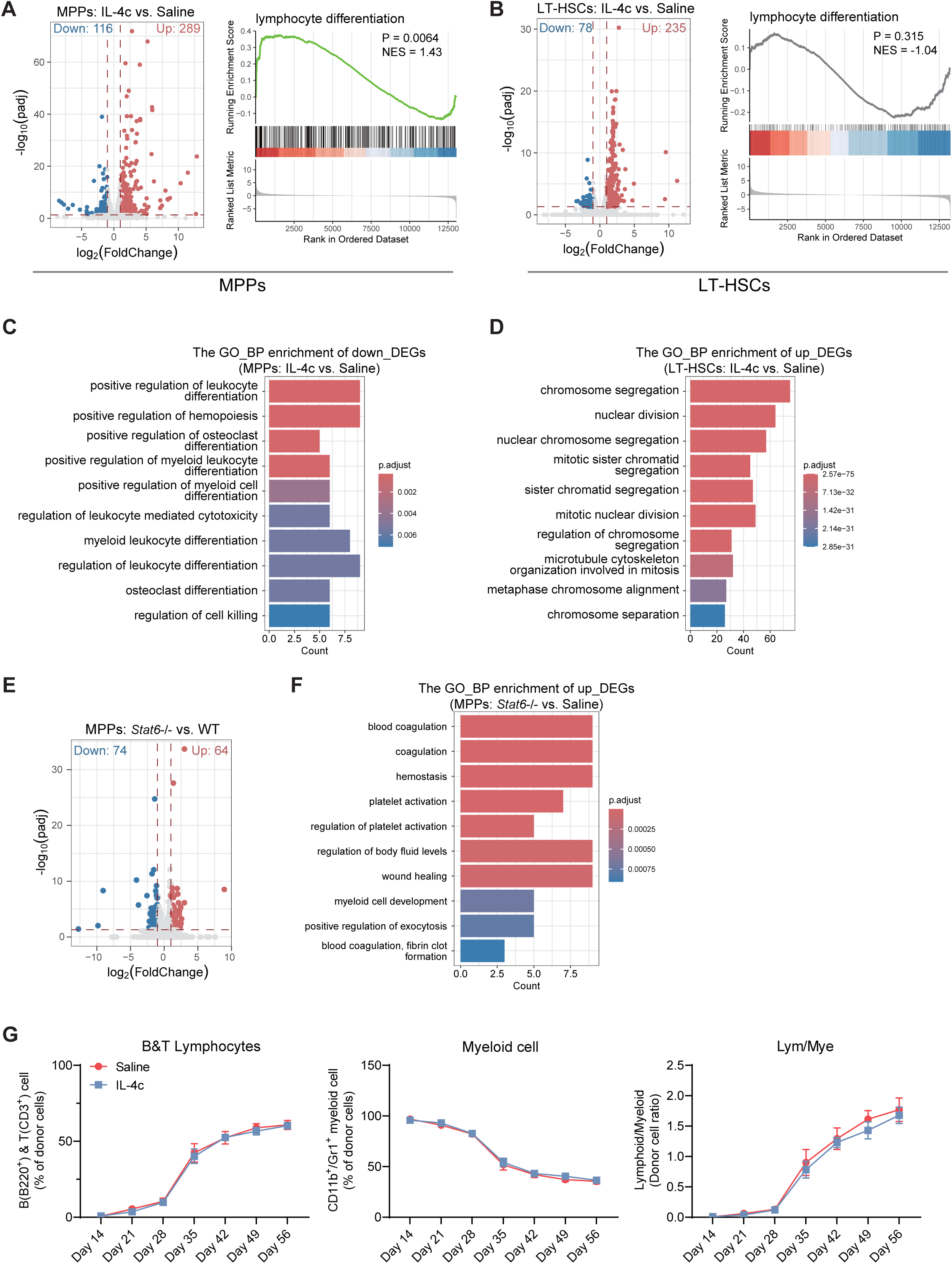
Transcriptomic analysis of MPPs and LT-HSCs after IL-4 treatment (Related to Figure 3) **A-B**, Volcano plot (left) showing the DEGs and GSEA enrichment (right) showing the “lymphoid differentiation” pathways in MPPs (**A**) or LT-HSCs (**B**) from IL-4c-treated versus saline-treated mice. **C-D**, GO pathway enrichment analyses of genes significantly downregulated in MPPs (**C,** log_2_FoldChange < –1, Padj<0.05) and upregulated in LT-HSCs (**D,** log_2_FoldChange > 1, Padj<0.05) from IL-4c-treated versus saline-treated mice. **E**, Volcano plot showing the DEGs in MPPs from *Stat6*^−/−^ versus WT mice. **F**. GO pathway enrichment analyses of genes significantly upregulated in MPPs from *Stat6*^−/−^ mice versus WT mice. (log_2_FoldChange > 1, Padj<0.05). **G**, Transplantation of LT-HSCs from IL-4c treated or untreated mice to lethally irradiated recipients: Donor-derived lymphoid cells (**G**, **left**), myeloid cells (**G**, **middle**) and lymphoid-to-myeloid ratio (**G**, **right**) in peripheral blood at the indicated days post-transplantation (n=5).

**Figure S6.**
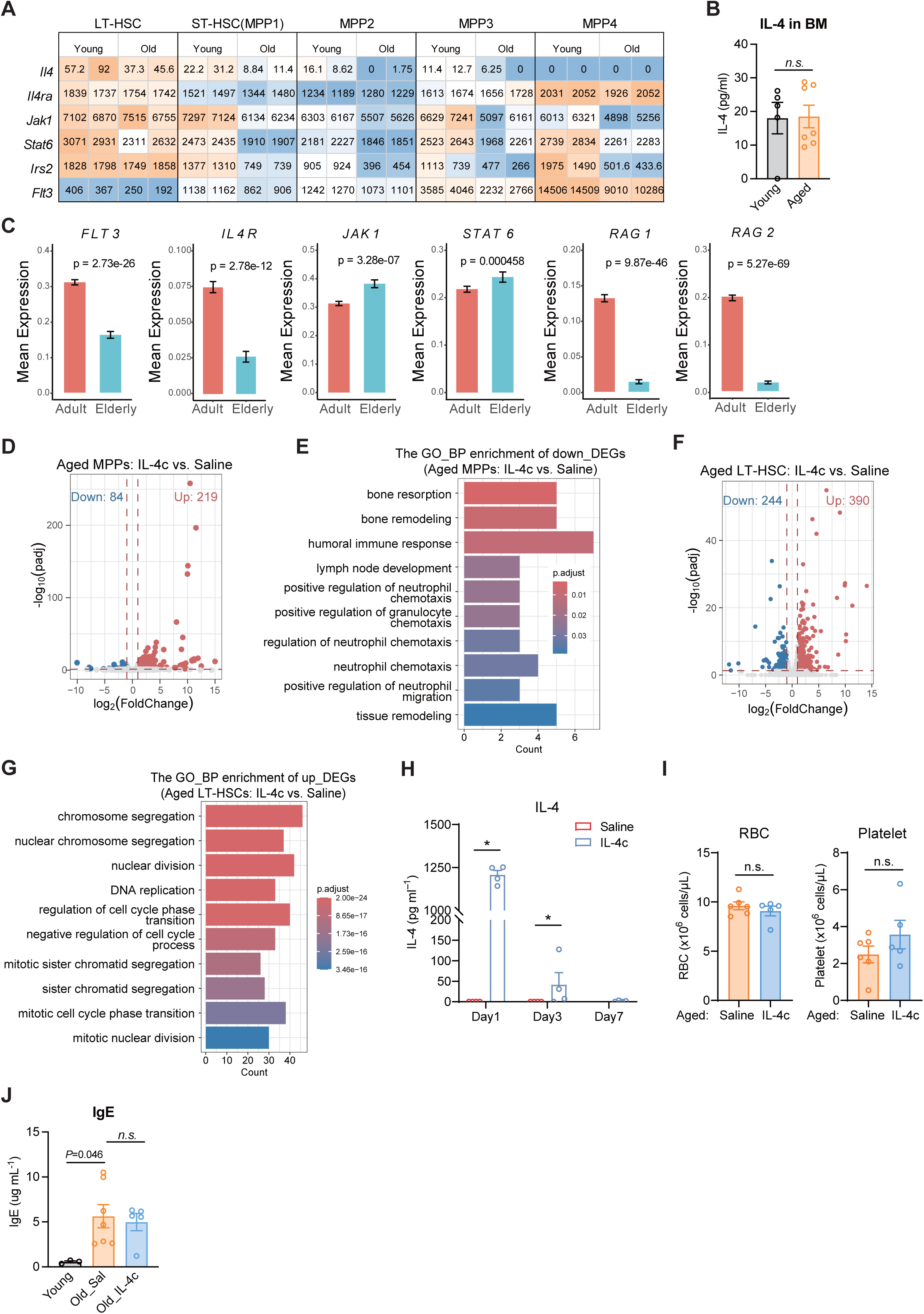
IL-4 effects on hematopoiesis in aged mice (Related to Figure 6) **A**, Heatmap of relative RNA expression of IL-4 downstream signaling molecules and *Flt3* in LT-HSC, ST-HSC and different MPPs from both young and aged mice. **B**, IL-4 level in young and aged BM fluid samples (n=5-7). **C**, IL-4 downstream signaling gene expression in human HSPCs from adult (25–53 years) and elderly (62–77 years) donors. **D**, Volcano plot showing the DEGs in MPPs from IL-4c-treated versus saline-treated aged mice. **E**, GO pathway enrichment analyses of genes significantly downregulated in MPPs from IL-4c-treated versus saline-treated aged mice (log_2_FoldChange < –1, Padj<0.05). **F**, Volcano plot showing the DEGs in LT-HSCs from IL-4c-treated versus saline-treated aged mice. **G**, GO pathway enrichment analyses of genes significantly upregulated in LT-HSCs from IL-4c-treated versus saline-treated aged mice (log_2_FoldChange > 1, Padj<0.05). **H**, IL-4 levels in serum following IL-4c injection, measured at indicated time points (n=4). **I**, Complete blood counting analysis of red blood cell (RBC) and platelet from aged mice with or without long-term IL-4c injections (n=5-6). **J**, IgE level in serum after long-term IL-4 treatment in aged mice (n=3-7). All data represent means ± s.e.m. Statistical significance was determined by one-way ANOVA with Tukey’s multiple-comparisons test (**J**) or unpaired two-tailed Student’s t-test (**B-C** and **H-I**).

**Figure S7.**
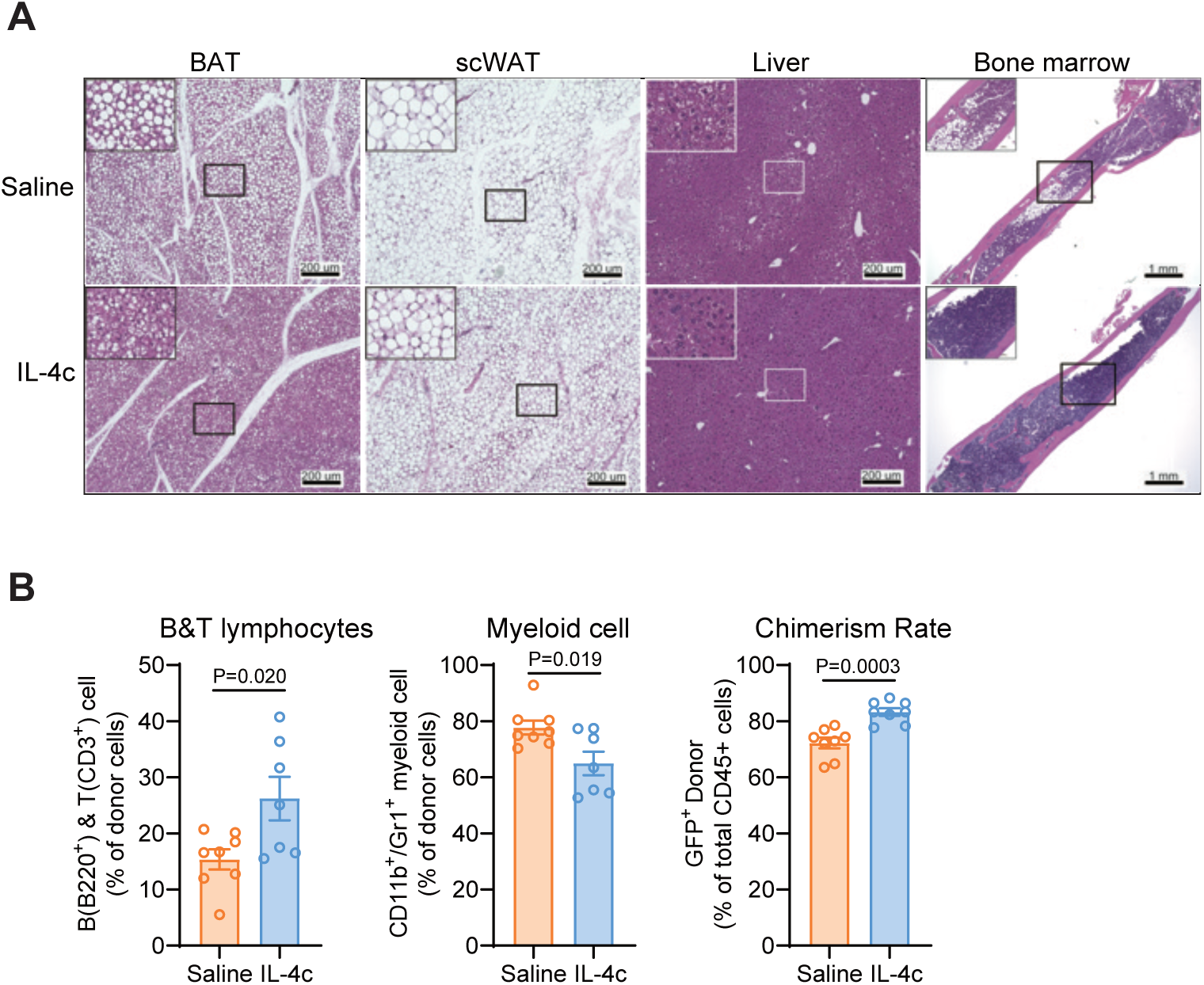
IL-4 treatment mitigates aging-associated decline in tissue function (Related to Figure 7) **A**, Representative images of hematoxylin-and-eosin (H&E) –stained sections of BAT (Brown adipose tissue), scWAT (Subcutaneous white adipose tissue), liver and bone marrow of saline or IL-4c treated aged mice. **B**, Donor-derived B and T lymphocytes (left), myeloid cells (middle) and chimerism rate (right) in peripheral blood one-month post-transplantation in aged mice (n=7-8). All data represent means ± s.e.m. Statistical significance was determined by unpaired two-tailed Student’s t-test (**B**).

